# PP4 inhibition sensitizes ovarian cancer to NK cell-mediated cytotoxicity via STAT1 activation and inflammatory signaling

**DOI:** 10.1101/2022.08.25.505192

**Authors:** Remya Raja, Christopher Wu, Thomas E Rubino, Emma Utagawa, Paul Magtibay, Kristina A. Butler, Marion Curtis

## Abstract

**Background:** Increased infiltration of T cells into ovarian tumors has been repeatedly shown to be predictive of enhanced patient survival. However, despite the evidence of an active immune response in ovarian cancer (OC), the frequency of responses to immune checkpoint blockade (ICB) therapy in OC is much lower than other cancer types. Recent studies have highlighted that deficiencies in the DNA damage response (DDR) can drive increased genomic instability and tumor immunogenicity, which leads to enhanced responses to ICB. Protein phosphatase 4 (PP4) is a critical regulator of the DDR; however, its potential role in anti-tumor immunity is currently unknown.

**Results:** Our results show that the PP4 inhibitor, fostriecin, combined with carboplatin leads to increased carboplatin sensitivity, DNA damage, and micronuclei formation. Using multiple ovarian cancer cell lines, we show that PP4 inhibition or *PPP4C* knockdown combined with carboplatin triggers inflammatory signaling via NFκB and STAT1 activation. This resulted in increased expression of the pro-inflammatory cytokines and chemokines: *CCL5, CXCL10*, and *IL6*. In addition, *IFNB1* expression was increased suggesting activation of the type I interferon response. Conditioned media from OC cells treated with the combination of PP4 inhibitor and carboplatin significantly increased migration of both CD8 T cell and NK cells over carboplatin treatment alone. Knockdown of STING in OC cells significantly abrogated the increase in CD8 T cell migration induced by PP4 inhibition. Co-culture of NK-92 cells and OC cells with *PPP4C* or *PPP4R3B* knockdown resulted in strong induction of NK cell activation as measured by IFN-γ levels. Further, we also observed increased degranulation and NK cell-mediated cytotoxicity against OC cells with *PPP4C* or *PPP4R3B* knockdown.

**Conclusions:** Our work has identified a role for PP4 inhibition in promoting inflammatory signaling and enhanced immune cell effector function. These findings support the further investigation of PP4 inhibitors to enhance chemo-immunotherapy for ovarian cancer treatment.

## BACKGROUND

High-grade serous ovarian cancer (OC) remains one of the most frequent causes for cancer-related death among women in the US [1]. Frontline treatment strategies for ovarian cancer patients include cytoreductive surgery combined with taxane or platinum-based chemotherapy. However, the emergence of platinum resistance is frequent, and the overall 5-year survival rate remains near 30% [2]. Immune checkpoint blockade (ICB) has been shown to induce durable responses in multiple cancer types. However, the benefit of ICB remains largely restricted to a minority of patients [3]. Multiple reports have confirmed that increased tumor-infiltrating T cells is correlated with improved clinical outcomes in OC [4, 5]. Despite these findings, ICB has produced minimal clinical benefit for ovarian cancer patients and as a result there are no FDA approved immunotherapies for OC [6]. These results underscore the need for the development of new rationally designed therapeutic approaches to improve outcomes for patients with OC.

Recent studies have shown that defects in DNA damage response (DDR) pathways can lead to increased genomic instability and tumor immunogenicity, which leads to enhanced responses to checkpoint inhibitor therapy [7, 8]. Homologous recombination (HR) defects are prevalent in OC and the introduction of poly (ADP-ribose) polymerase (PARP) inhibitors have opened new therapeutic avenues.

Protein phosphatase 4 (PP4) is a multi-subunit Ser/Thr phosphatase complex that is involved in diverse cellular pathways including DNA damage repair, cell cycle progression, and apoptosis [9]. PP4 plays a central role in the response to DNA damage as it directly dephosphorylates many proteins that play crucial roles in the DDR, including γH2AX, Kap-1/TRIM28, RPA2, and 53BP1 [10-13]. Dephosphorylation of these critical substrates is necessary for resolution of DNA damage repair. We previously discovered that the cancer/testis antigen, CT45, directly binds to the PP4 complex and functions as an endogenous inhibitor of PP4 activity leading to enhanced platinum sensitivity and DNA damage. In addition, patients whose tumors have increased levels of the catalytic subunit, PP4C, were found to have reduced overall survival [14]. However, the role of PP4 in anti-tumor immunity is currently not known. In this study, we used a combination of the PP4 inhibitor, fostriecin, and transient PP4 knockdown to interrogate the role of PP4 in chemosensitivity and in immune effector function in OC.

## METHODS

### Cell lines

High-grade serous ovarian cancer cell lines NIH-OVCAR3, OVCAR4 and OVCAR8 were purchased from NCI-Frederick Cancer DCTD Tumor/Cell Line repository and ID8 was purchased from Millipore. The ovarian cancer cell lines, NIH-OVCAR3, OVCAR4, and OVCAR8 were cultured in RPMI with 10% FBS and 1% penicillin/streptomycin. The mouse ovarian cancer cell line, ID8, was cultured in DMEM with 4% FBS, 1X Insulin-Transferrin-Selenite and 1% penicillin/streptomycin. NK-92 cell line was a kind gift from Abhinav Acharya, Arizona State University, USA. OVCAR8-DRGFP cell line was a kind gift from Dr. Larry Karnitz, Mayo Clinic, Rochester, USA. The cells were grown in complete RPMI media with 30 µg/ml puromycin. All cell lines were tested for mycoplasma and authenticated using a commercial service (CellCheck, IDEXX Bioresearch).

### p53 CRISPR knockout cell generation

A p53 CRISPR knockout clone was generated from ID8 (ID8-p53KO) and was cultured as the parental cell line (ID8-p53 WT). Three crRNA targeting the DNA binding domain of p53 was selected as described previously [15]. Pre-designed guides with the highest on-target and cutting efficiency scores were purchased from Integrated DNA Technologies in their proprietary Alt-R format **(Supplementary table 1)**. crRNA-tracrRNA duplexes were prepared by annealing equimolar concentrations of AltR crRNA with Alt-tracrRNA-ATTO 550 at 95°C for 5 min in a thermocycler and allowed it to slowly cool to room temperature. In a PCR strip, three crRNA-tracrRNA duplexes were mixed Recombinant Cas9 V3 protein (#1081058, IDT) and incubated for 20 min. ID8 cells were electroporated (BTX ECM 830, Harvard Apparatus) with ribonucleoprotein complex and electroporation enhancer (#1075915, IDT). The gene edits in single clones were verified by Sanger Sequencing and p53 protein expression was confirmed using western blot.

### RNA-mediated gene knockdown

Human and mouse siGenome SMARTpool *PPP4C* siRNA, human *PPP4R3B*, ON-TARGETplus SMARTpool human *STING1* siRNA and non-targeting control pool #2 were purchased from Horizon Discovery Biosciences. Transfection with siRNAs were performed with Lipofectamine 3000 as per manufacturer’s protocol.

### Bioinformatics analysis

Genome profiling of PP4 complex members namely *PPP4C, PPP4R3A, PPP4R3B* and *PPP4R2* across different cancer types was done using the pan-cancer TCGA data available at cBioportal (https://www.cbioportal.org/) [16]. The Oncoprint for PP4 subunits in ovarian cancer (TCGA, pan-cancer atlas) was also created using the relevant module at cBioportal. RNAseq expression values (TPM) were obtained for *PPP4C, PPP4R3A,PPP4R3B* and *PPP4R2* from TCGA and GTex using the UCSC Cancer Browser (https://xena.ucsc.edu/welcome-to-ucsc-xena/) [17]. From Depmap database (https://depmap.org/portal/), RNA level expression values and proteomic scores for PP4C were obtained for relevant ovarian cancer cells and plotted [18]. The TIMER 2.0 (https://timer.cistrome.org/) gene module was used to analyze the correlation between the tumor-specific gene expression of PP4 subunits and immune cell infiltration, particularly for NK and NKT cell populations. The purity corrected ovarian cancer dataset (n=303) was analyzed using benchmarked CIBERSORT and xCELL deconvolution tools [19, 20].

### Incucyte live-cell imaging and cell viability analysis

For cell viability analysis with fostriecin, the ID8 p53-WT cell line was seeded in a 96-well plate. At 24h, cells were treated with fostriecin (1nM), the control cells were left untreated. The next day, control and fostriecin pre-treated cells were treated with varying concentrations of carboplatin (1µM-1mM). For live cell imaging and cell proliferation, IncuCyte images were taken every 4 hours and cell confluence was recorded. For siRNA experiments, both ID8-p53WT and p53KO cells were transfected with *PPP4C* siRNA followed by carboplatin treatment for 96h. Cell viability was measured by CellTitre 96 Aqueous Non-Radioactive Cell Proliferation Assay (#G5421, Promega) as per manufacturer’s instructions.

### Clonogenic survival assay

Cells were treated with different carboplatin concentrations (as indicated in the figure legends) for 96 hrs. 2000 cells per 6-well were then plated for 7 days in drug-free medium. Grown colonies were fixed with 1% formaldehyde for 10 min at room temperature and stained with 40% methanol and 0.05% Crystal Violet for 20 min. The colonies were counted using Colony Area ImageJ plugin and plotted as percent survival of control [21].

### Immunoblotting

Western blot was performed as described previously [14]. Ovarian cancer cells were pre-treated with fostriecin (1nM) on day 1, post-seeding. The cells were treated with carboplatin at indicated doses ± fostriecin (1nM) on day 2. On day 5, cells were collected in RIPA buffer and lysates prepared for SDS-PAGE and immunoblot analysis. The list of antibodies used in the study are listed in Supplementary Table 2.

### Phosphatase activity assay

Cellular protein phosphatase 4 (PP4) activity in ovarian cancer cells was performed as described previously with minor modifications [14]. Immunoprecipitation was carried out from lysates (1mg) using anti-PPP4C antibody or Control IgG, followed by incubation with Protein-A agarose beads (#22811, Thermo Scientific) overnight at 4°C. The washed beads were re-suspended in 100 µl assay buffer (30 mM of HEPES, 0.1 mg/mL of BSA, 0.1 mM of MnCl2, 1mM of sodium ascorbate, 1 mM of DTT, 0.01% Triton X-100). ± fostriecin (1nM) and incubated at room temperature for 30 min. PP4 activity was assayed by incubating the washed beads with 100 µM DiFMUP substrate (#D6567, Invitrogen) and fluorescence measured at 450 nM after 60 minutes using FlexStation 3 (Molecular Devices). PP4 activity is shown as percentage of untreated control.

### Immunofluorescence and micronuclei analysis

OVCAR3, OVCAR4, OVCAR8 and ID8-p53KO cells were seeded in a 96-well plate and treated ± fostriecin (1nM), followed by carboplatin at indicated doses: OVCAR3 and 4 (1µM), OVCAR8 (2.5 µM) and ID8-p53KO (10 µM). At 96hr, cells were washed with 1X PBS, followed by fixation for 10min at room temperature with 4% paraformaldehyde. The cells were stained with Hoechst 33342 (#H3570, Invitrogen) for 10 min and micronuclei were imaged using Image Xpress (Molecular Devices). A minimum of 45-50 sites were imaged per conditions and represented as violin plots. Immunofluorescence analysis with γ-H2AX (S139), (#MA1-2022, Thermo Fischer Scientific) and FANCD2 (#ab108928, Abcam) in OVCAR3 was performed as described previously [14].

### Cellular thermal shift assay (CETSA)

The assay was performed as described previously with minor modifications [22, 23]. Briefly, OVCAR4 and OVCAR3 cells were collected in PBS with 1mM sodium ascorbate, incubated ± fostriecin (1nM) for 30 min and subjected to increasing temperatures. Cells were lysed and immunoblotting was done for PP4C. The densitometry on immunoblots was performed with Image J (NIH), and relative band intensities were quantified as percentage of non-denatured protein.

### qRT-PCR

The ovarian cancer cells were treated with fostriecin (1nM) on day 1, followed by carboplatin treatment at indicated doses ± fostriecin on day 2. The cells were harvested on day 5 post carboplatin treatment for quantitative PCR. For siRNA experiments, cells were transfected with *PPP4C* or control siRNA on day 1, followed by carboplatin treatment on day 2. The cells were harvested on day 5. The primers used for qRT-PCR are listed in Supplemental Table 3.

### DR-GFP homologous recombination (HR) assay

HR assays were performed using OVCAR8-DR-GFP cells. The experiments were performed as described previously with minor modifications [24]. For siRNA experiments, the cells were transfected twice. On day 1, they were transfected with *PPP4C* or control siRNA only. On day 2, cells were transfected with 20□μg pCßASceI plasmid (encoding I-SceI) with empty vector (pcDNA3.1) and mCherry (pCDNA3.1). GFP and mCherry fluorescence were assessed by flow cytometry on day 5. OVCAR8-DR GFP cells were transfected with pCßASceI plasmid (encoding I-SceI) with empty vector (pcDNA3.1) and mCherry (pCDNA3.1) on Day 1, followed by fostriecin (1nM) treatment daily till flow analysis on day 3 post-transfection.

### CD8 T cell and NK cell migration

ID8-p53KO or OVCAR3 cells were treated with fostriecin (1nM) on day 1, followed by carboplatin (10µM) ± fostriecin (1nM) on day 2. The media was changed to basal RPMI-1640+0.5% BSA after 72 h of carboplatin treatment. The conditioned media was collected after 48 hours. OT-I splenocytes were isolated and stimulated with OVA in presence of IL-2. On day 3 following OVA stimulation, CD8 T cells were enriched using mouse CD8a T cell positive selection kit (#18953, STEMCELL) according to manufacturer’s instruction. Mouse CD8 T or NK-92 (1e5) cells were placed in basal media in the upper chamber of 5-micron filters, conditioned media corresponding to various treatments from ID8 and OVCAR3 was placed in the lower chamber respectively [25, 26]. The migrated cells were collected at 4h and quantified with Cell Titre-Glo 2.0 reagent (#G9241, Promega). For inhibitor experiments, CD8 T cells or NK-92 cells were incubated with either STATTIC (2 µM) or Trametinib (7.5 µM) or NSC23766 (10 µM) for 15 minutes at 37°C, prior to adding to the upper chamber.

### Intracellular staining

OT-I splenocytes were stimulated with OVA (30ng/ml) ± fostriecin at 1 and 10nM. At 72h, cells were restimulated with OVA peptide in presence of Golgiplug and incubated for 4 h at 37°C in a 5% CO2 incubator. The cells were surface markers (anti-mouse-CD3, -CD8, live/dead stain). Cells were fixed, permeabilized and stained for cytokines using an anti-IFN-γ and TNF-α antibody. For cell co-culture experiments, OVCAR8 cells were treated with fostriecin or transfected with either *PPP4C* or control or *PPP4R3B* siRNA, followed by carboplatin treatment (2.5 µM). On Day 5, cells were trypsinized and seeded on to a flat bottom 12 well plate. After 6 h, NK-92 cells were added in 1:1 (tumor: NK cell) ratio. At the end of 14 h co-culture, NK-92 cells were collected and restimulated with cell stimulation cocktail (#TNB-4975-UL100, Tonbo Biosciences) for 4 h and intracellular staining was performed. All antibodies used in flow cytometry are provided in Supplementary table 4. Flow data analysis was performed with FlowJo (Version 10.8.0)

### CD107a mobilization and NK cell cytotoxicity assay

OVCAR8 cells were transfected with respective siRNAs and treated with carboplatin at 2.5µM. On day 5, cells were washed, trypsinized and mixed with NK-92 in 1:1ratio (target: effector) in a U-bottom 96 well plate. CD107a antibody was added at the beginning of the experiment and cells were co-cultured for 3h in the presence of Golgi Plug and Monensin. At the end of 3h, cells were stained with cell surface markers and analyzed by flowcytometry. For cytotoxicity assays, OVCAR8 cells were treated as described previously and on day 5, cells were mixed with NK-92 cells in 1:1 ratio (target: effector) in a U-bottom 96 well plate. At the end of 3 h incubation, cells were stained with anti-Annexin V and PI.

### Statistical Analysis

All statistical analyses were performed with GraphPad Prism (GraphPad) software V.9. For experiments with one comparison, unpaired two-tailed t-test was used. The mean and the standard error of the mean (SEM) or standard deviation (SD) are reported for all graphs. For experiments with more than one comparison, One-way ANOVA with Tukey’s multiple comparisons post-test was used. Before applying ANOVA, we first tested whether the variation was similar among the groups using the Bartlett’s test. Differences were considered significant if p<0.05. No blinding or randomization was done during data acquisition or assessment of outcome and sample sizes were determined based on previous experience with the individual experiment.

## RESULTS

### Overexpression of PP4 subunits in ovarian cancer

The PP4 holoenzyme is comprised of the catalytic subunit (PP4C) and multiple different regulatory subunits: PP4R1, PP4R2, PPP4R3β and PPP4R3α. The catalytic subunit forms distinct complexes with different regulatory subunits that have been shown to influence both PP4 enzyme activity and substrate selectivity [11]. We previously identified CT45 as an endogenous inhibitor of PP4 complex, that physically interacts with the PP4C-PPP4R2-PPP4R3α/β complex leading to enhanced carboplatin sensitivity in OC [14]. To better understand the role of PP4 in cancer we investigated the genome level alterations of *PPP4C, PPP4R3A, PPP4R3B* and *PPP4R2* subunits using the pan-cancer TCGA data in cBioportal (n=10953). We found that the PP4 subunit genes are predominantly amplified in roughly 6% of OC tumors **(Figure 1A, Supplementary Figure 1A)**. Similar to OC, invasive breast carcinoma, bladder urothelial carcinoma, and seminoma also showed significant amplification of PP4 subunit genes **(Figure 1A)**. In addition, we observed significant concordance between mRNA levels and amplification of *PPP4C* in the pan-cancer TCGA data **(Figure 1B)** as well as for the regulatory subunits **(Supplementary Figure 1B)**. To assess the utility of targeting the PP4 complex in OC we next evaluated the expression levels of PP4 subunits in the OC TCGA dataset. Using RNAseq data, we found that *PPP4C* is significantly overexpressed in OC as compared to normal tissue **(Figure 1C)**, while the expression levels of the regulatory subunits were found to be variable **(Figure 1C)**. Increased mRNA expression was noted for *PPP4R3B* and *PPP4R2*. In contrast, *PPP4R3A* was found to be higher in normal tissue **(Figure 1C)**. At the protein level, PP4C was found to be robustly expressed across a panel of representative HGSOC cell lines **(Figure 1D, panel i)**. We obtained *PPP4C* RNA expression values and proteomic scores for relevant HGSOC cell lines from the Depmap portal. Both RNA level expression and proteome scores were consistent with the immunoblot data **(Supplementary Figure 1C)**. Interestingly, we noted variable expression of the PPP4R3β regulatory subunit across the OC cell lines, with TYK-nu and 59M showing the lowest expression **(Figure 1D, panel i)**. A similar trend was also observed with the other two regulatory subunits, PPP4R3α and PPP4R2 **(Figure 1D, panel ii)**. Taken together, these data suggest that PP4C is frequently upregulated in OC.

**Figure 1:**
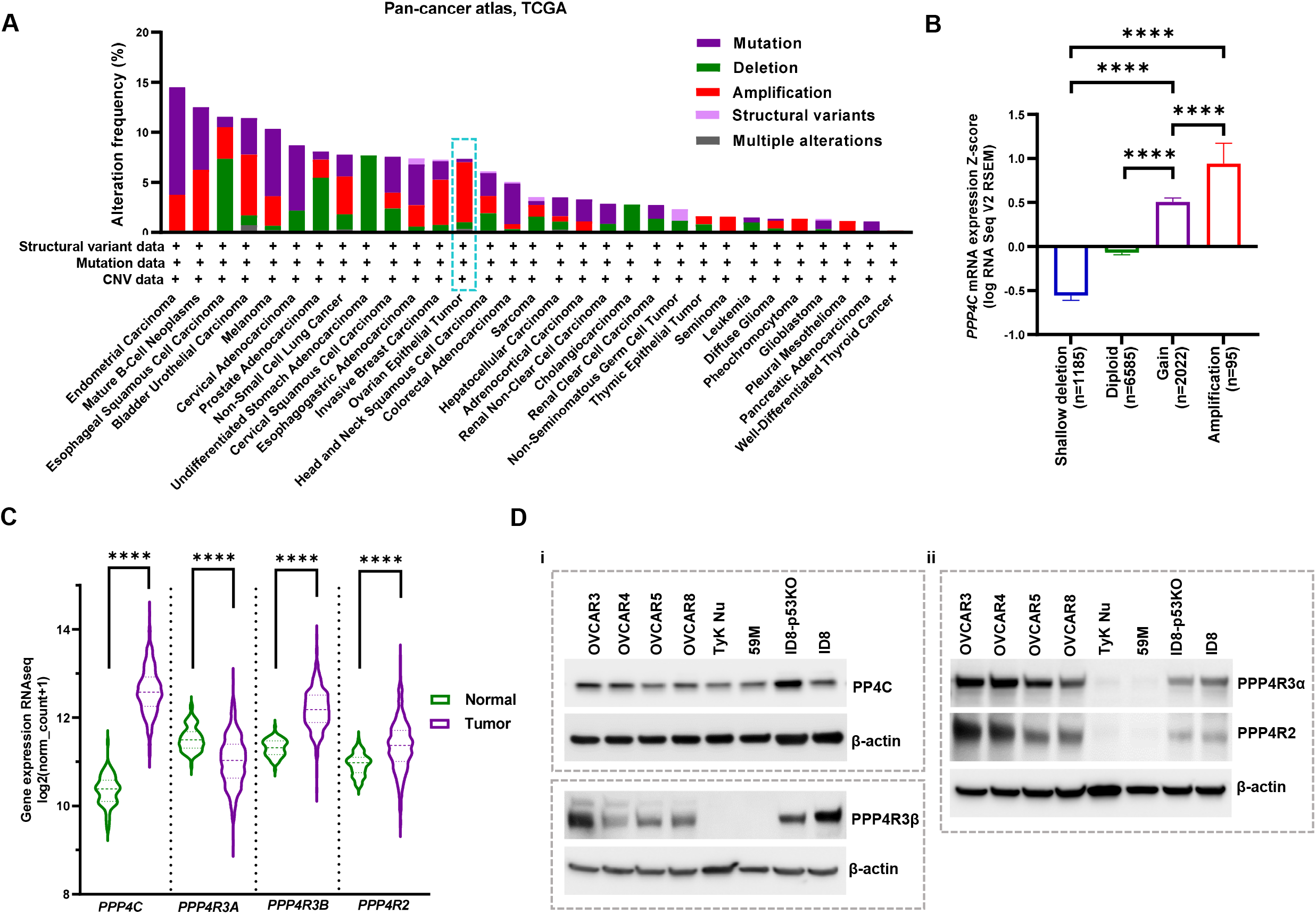
PP4 subunits are amplified in ovarian cancer. (A) Genomic profiling of PP4 across human cancers using TCGA data from cBioPortal. (B) *PPP4C* mRNA expression versus copy number from pan-cancer TCGA data is plotted as a bar graph, ****p<0.0001. (C) Comparison between transcript level expression of PP4 subunits (TPM) compared between TCGA ovarian tumor cohort (n=418) and normal (GTex) (n= 88), ****p<0.0001.(D) i, ii, Western blot showing expression of PP4 subunits across a panel of ovarian cancer cells.

### Gene silencing or pharmacological inhibition of PP4C enhances sensitivity to carboplatin in ovarian cancer

Given the known role of PP4 in the DDR, we next sought to ascertain the effect of PP4 inhibition on sensitivity to carboplatin, which is commonly used clinically in OC. *TP53* mutations are prevalent and are found in ∼90% of all ovarian cancers. As ID8 is a p53 wild-type cell line, we generated a p53 knockout ID8 cell line model using CRISPR. The gene edits in the targeted region were verified by Sanger sequencing and loss of p53 expression in the knockout clone was verified by western blot (**Supplementary Figure 2 A-C**). *PPP4C* knockdown with siRNA in ID8-p53WT and ID8-p53KO showed that loss of PP4C expression led to increased sensitivity to carboplatin in both the cell lines (**Figure 2A**). PP4C expression after siRNA transfection was verified by western blot (**Figure 2A, inset**). Similar to the siRNA results, treatment with pharmacological inhibitor of PP4, fostriecin, at nanomolar levels also resulted in enhanced sensitivity to carboplatin in ID8 mouse cell line (**Figure 2B**). Decreased colony-forming capacity was seen upon treatment with fostriecin and carboplatin in ID8-p53KO and the human OC cell lines, OVCAR3, OVCAR4, and OVCAR8 compared to carboplatin treatment alone (**Figure 2C i& ii**). These findings demonstrate that suppressing PP4 activity can lead to increased carboplatin sensitivity in OC. Additionally, we confirmed that PP4 phosphatase activity was inhibited by fostriecin in OVCAR3 and OVCAR4 human OC cell lines (**Supplementary Figure 3**). PP4C was confirmed to be a direct target of fostriecin using a Cellular Thermal Shift Assay (CETSA) in OC cell lines (**Figure 2D**).

**Figure 2:**
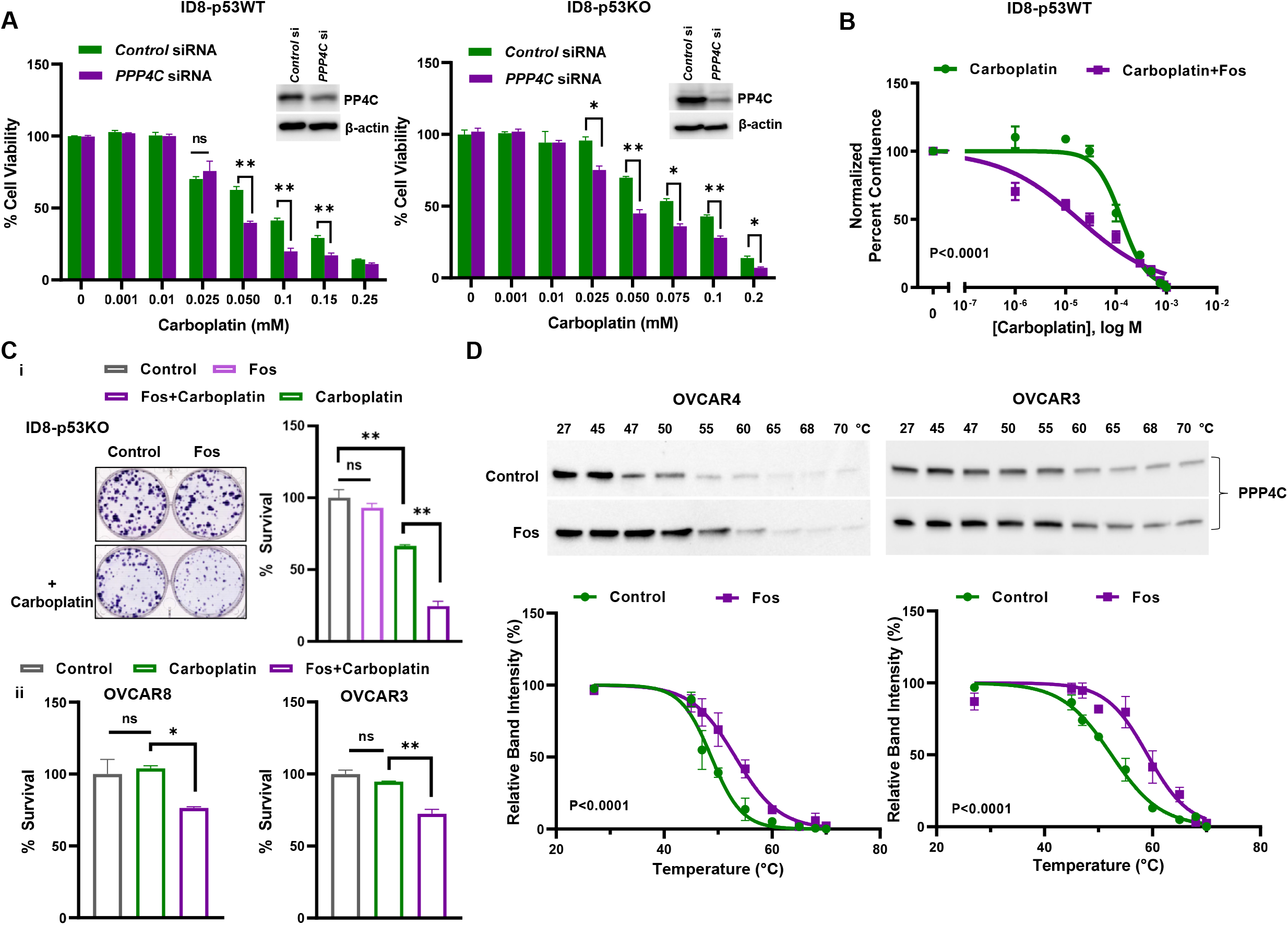
PP4 inhibition or knockdown sensitizes OC cell lines to carboplatin. (A) Mouse ovarian cancer cell line, ID8 (ID8-p53WT) and isogenic p53 KO cell line (ID8-p53KO) cells were transfected with control or *PPP4C* siRNA. Cell viability was measured by MTS assay at 96 h, results represent three replicates per experiment group, *p < 0.05, p**<0.01. In parallel, PP4C knockdown following siRNA transfection was verified by western blot (inset). (B) ID8-p53WT cells were treated ± Fostriecin (Fos) (1nM) for 24 h, followed by carboplatin treatment at indicated doses. Cell proliferation was monitored using IncuCyte S3 and the difference in mean cell confluence across treatment groups (n=6) is plotted, *p<0.0001. (C) i, Clonogenic survival assay of ID8-p53KO cells treated ± Fos (1nM), followed by carboplatin (10µM) treatment. Representative images are shown adjacent to the graph. ii, Human ovarian cancer cell lines OVCAR3 and OVCAR8 cells were treated ± Fos (1nM), followed by carboplatin (1µM) and colony survival was determined. Mean values are shown from three independent experiments. Error bars show SEM for each group, *p<0.05, **p<0.01. (D) CETSA assay on OVCAR4 and OVCAR3 cells ± Fos (10nM). Data shown is an average of 3 biological replicates, *p<0.0001.

### Inhibition of PP4C enhances carboplatin induced DNA damage

Genomic instability is a hallmark of cancer. Deficiencies in the DNA damage response can lead to genomic instability and can make cancer cells more sensitive to chemotherapy. Replication stress caused by chemotherapeutics that induce double-strand breaks, such as carboplatin, can lead to micronuclei formation, a manifestation of genomic instability. We found that PP4 inhibition with fostriecin combined with carboplatin led to increased micronuclei formation in multiple OC cell lines **(Figure 3A)**. As PP4 has a known role in DNA damage repair, we sought to determine if fostriecin treatment affected H2AX (S139) phosphorylation (γ-H2AX), which is a well-reported substrate for PP4 phosphatase [11]. Our results show that fostriecin treatment in combination with carboplatin enhanced overall γ-H2AX levels in OC cells. However, we did not observe any significant changes in γ-H2AX with fostriecin treatment alone **(Supplementary Figure 4A)**. γ-H2AX foci formation, was also increased in OVCAR3 following treatment with the combination of fostriecin and carboplatin as compared to carboplatin or fostriecin alone (**Figure 3B**). Since carboplatin is known to induce inter-strand crosslinks (ICLs), we next investigated whether FANCD2 foci, a critical sensor of ICLs [27], were altered by PP4 inhibition. Consistent with literature, we found increased FANCD2 foci in response to carboplatin treatment; however, fostriecin treatment did not increase the number of FANCD2 foci per cell **(Figure 3B)**. Chowdhury *et al*. had previously reported an increase in basal γ-H2AX upon loss of PP4C. Similar to their findings, we also observed increased overall γ-H2AX levels as measured by intensity in fostriecin and carboplatin treated OVCAR3 cells. However, no difference was observed in FANCD2 basal expression with addition of fostriecin to carboplatin treated cells **(Supplementary Figure 4B**). Because PP4 is known to play a role in homologous recombination (HR) [10], we next determined whether loss of PP4 affects HR activity in OC cells. *PPP4C* knockdown via siRNA and fostriecin treatment inhibited homologous recombination (HR) as shown using the HR reporter cell line, OVCAR8-DR-GFP **(Figure 3C)**. These data demonstrate that inhibition of PP4 leads to increased DNA damage and markers of genomic instability in OC cell lines following treatment with carboplatin.

**Figure 3:**
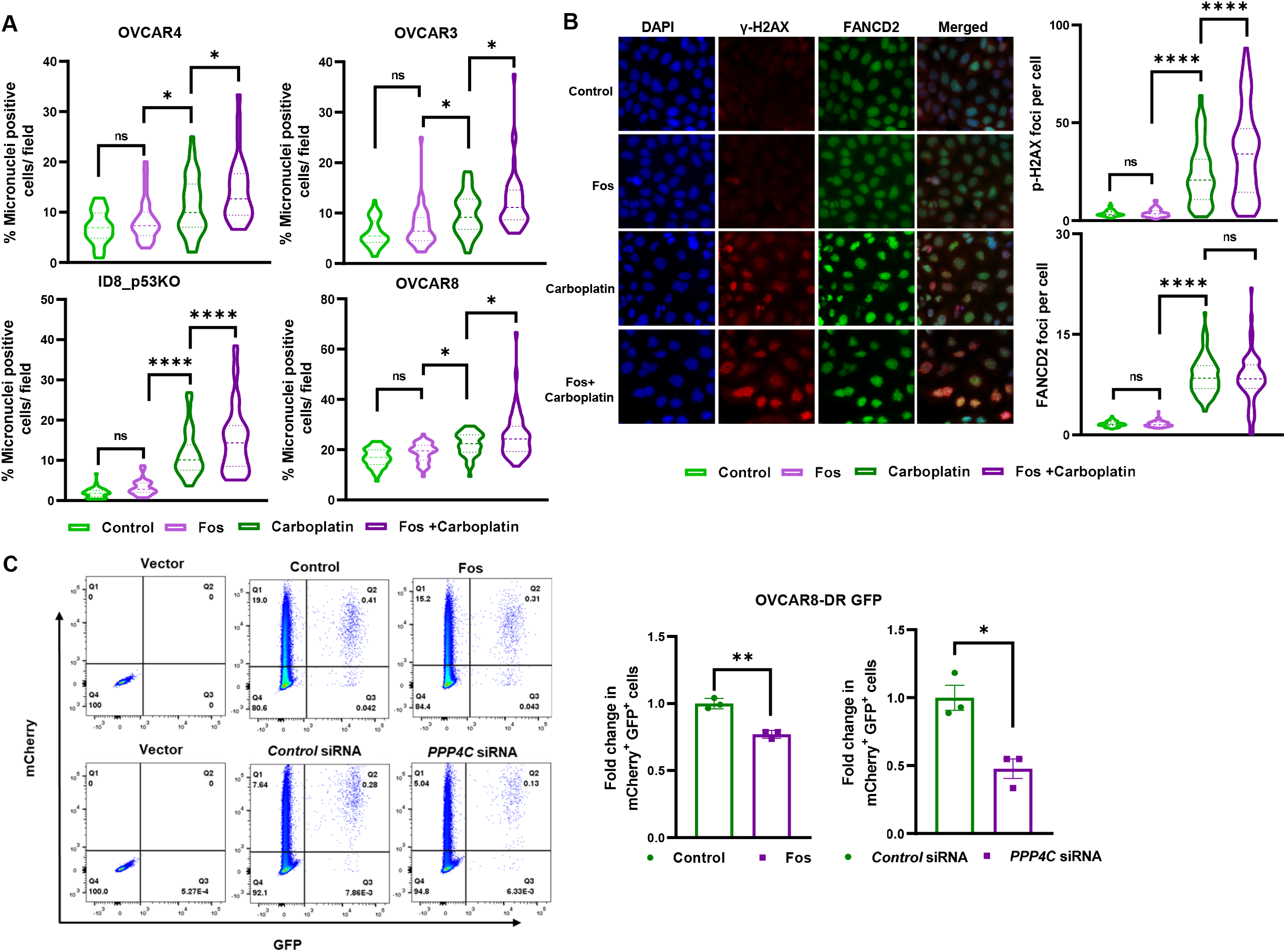
Pharmacological inhibition of PP4 augments carboplatin-induced DNA damage. (A) OC cell lines were treated with Fos (1nM) followed by carboplatin at indicated doses, OVCAR4 and OVCAR3 (1 µM), ID8-p53KO (10 µM) and OVCAR8 (2.5 µM). At 96 h, micronuclei formed across conditions were imaged. The micronuclei count per 45-50 sites was averaged across conditions and represented as violin plots, *p<0.05, ****p<0.0001. (B) OVCAR3 cells were treated ± Fos (1nM), followed by carboplatin treatment at 2.5 µM for 96 h and co-immunostained with FANCD2 and γ-H2AX. Representative images of γ-H2AX and FANCD2 foci across treatment conditions are shown. Image-based quantitation of foci counts was done across 70-90 sites and are shown in violin plots. ****p<0.0001. (C) OVCAR-8-DR-GFP cells were transfected with pCβASceI plasmid and treated ± Fos (1nM) every day for 72 h or transfected with control or *PPP4C* siRNA. Cells were analyzed for GFP fluorescence and efficiencies in HR were normalized to either untreated controls or control siRNA transfected cells. Values are mean ±SEM. *p<0.05, **p<0.01.

### PP4 inhibition promotes inflammatory signaling in ovarian cancer

The presence of cytosolic DNA such as micronuclei can lead to activation of the cGAS-STING pathway, which can trigger pro-inflammatory signaling and synergize with immunotherapy to promote anti-tumor immunity [28]. Due to the increase in micronuclei observed with PP4 inhibition **(Figure 3A)**, we hypothesized that inflammatory signaling would be increased in OC cells upon loss of PP4 activity. Signal transducer and activator of transcription (STAT) 1 is a key mediator of interferon signaling and plays an important role in both innate and adaptive immunity [29]. STAT1 phosphorylation at Y701 is essential for its activation and nuclear translocation. Additional phosphorylation at S727 is required for full transcriptional activation of STAT1. We found that treatment with the combination of fostriecin and carboplatin led to increased STAT1 (Y701) phosphorylation in OC cell lines as compared to carboplatin treatment alone **(Figure 4A, Supplementary Figure 5A)**. Similar to fostriecin treatment, knockdown of *PPP4C* combined with carboplatin also resulted in increased STAT1 activation in OC cell lines **(Figure 4B & Supplementary Figure 5B)**. The loss of perinuclear envelope surrounding the micronuclei, can trigger pattern recognition receptor cyclic GMP-AMP synthase (cGAS), which in turn elicits stimulator of interferon genes (STING)-mediated type I interferon response resulting in STAT1 phosphorylation [30]. STING activates several transcription factors including IRF3 and NF-κB leading to the production of type I interferons and proinflammatory cytokines [31]. Consistent with literature, we observed significant upregulation of phospho-p65 in response to fostriecin and carboplatin in both OVCAR3 and OVCAR8 cells (**Figure 4C**). Canonical NF-kB was also found to be activated upon knockdown of *PPP4C* expression in these cell lines, which was further increased upon carboplatin treatment (**Figure 4D)**. As we observed increased phospho-p65 in response to PP4C loss, we next determined the levels of *IFNB1* and select proinflammatory chemokines and cytokines in OC cells. At the mRNA level, *IFNB1, CCL5, CXCL10*, and *IL6* were found to be increased with the combination treatment of carboplatin with fostriecin **(Figure 4E, Supplementary Figure 5C, left)** or *PPP4C* siRNA **(Figure 4F, Supplementary Figure 5C, right)**. Collectively, these data demonstrate that PP4 inhibition augments carboplatin-induced inflammatory signaling in OC cell lines.

**Figure 4:**
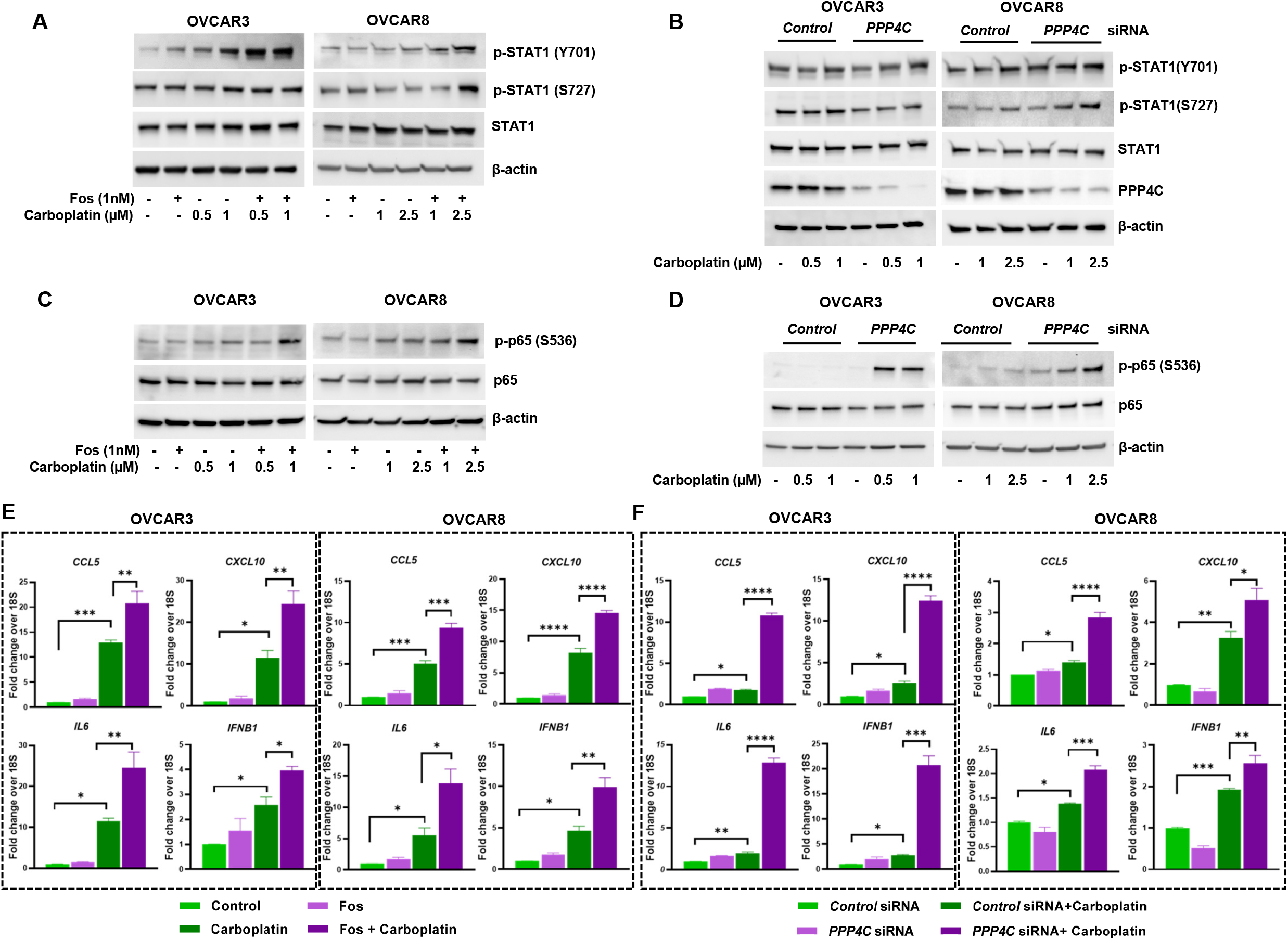
Loss of PP4 activity potentiates DNA damage-induced inflammatory signaling. (A) Representative western blot showing STAT1 phosphorylation at Y701 and S727 in OC cells treated ± Fos(1nM) followed by carboplatin treatment at indicated doses on Day 5. (B) STAT1 signaling shown following *PPP4C* siRNA transfection and carboplatin treatment on Day 5. Loss of PP4C expression is also shown. (C) Representative western blot showing p65 phosphorylation at S536 in HGSOC cells treated ± Fos(1nM) followed by carboplatin treatment at indicated doses on Day 5. (D) Immunoblots of p-p65 following PPP4C siRNA transfection and carboplatin treatment on Day 5. Actin is used as loading control. (E, F) Transcript levels of *CCL5, CXCL10, IL6* and *IFNB1* were measured in OVCAR3 and OVCAR8 cells ± Fos (E) or *PPP4C* siRNA transfection (F) and carboplatin treatment by quantitative, real-time PCR and relative fold change was calculated from 3 replicates. Values are mean ±SEM. *p<0.05, **p<0.01,***p<0.001 and ****p<0.0001.

### PP4 inhibition in ovarian cancer cells boosts immune cell migration via the STING pathway

Previous studies have shown that increased CD8 T cell infiltration correlated with improved prognosis in OC [4]. CCL5, CXCL9, and CXCL10 have been shown to enhance both T and NK cell infiltration into tumors [32, 33]. To determine whether proinflammatory cytokines and chemokines produced upon combination of carboplatin and PP4 inhibition influences immune cell migration, we collected conditioned media from both mouse and human OC cell lines following treatment with fostriecin and carboplatin. Mouse OT-I CD8 T cells or NK-92 cells were placed in a Boyden chamber with either the mouse or human OC conditioned media, respectively **(Figure 5A, schematic)**. Both mouse CD8 T cells and NK-92 cells displayed enhanced migration when exposed to conditioned media from OC cells treated with the combination of fostriecin and carboplatin **(Figure 5A)**. Additionally, we verified that direct fostriecin treatment had no effect on T cell viability and did not negatively influence ovalbumin-induced T cell activation as shown by intracellular cytokine staining for IFNγ and TNFα **(Supplementary Figure 6A)**.

**Figure 5:**
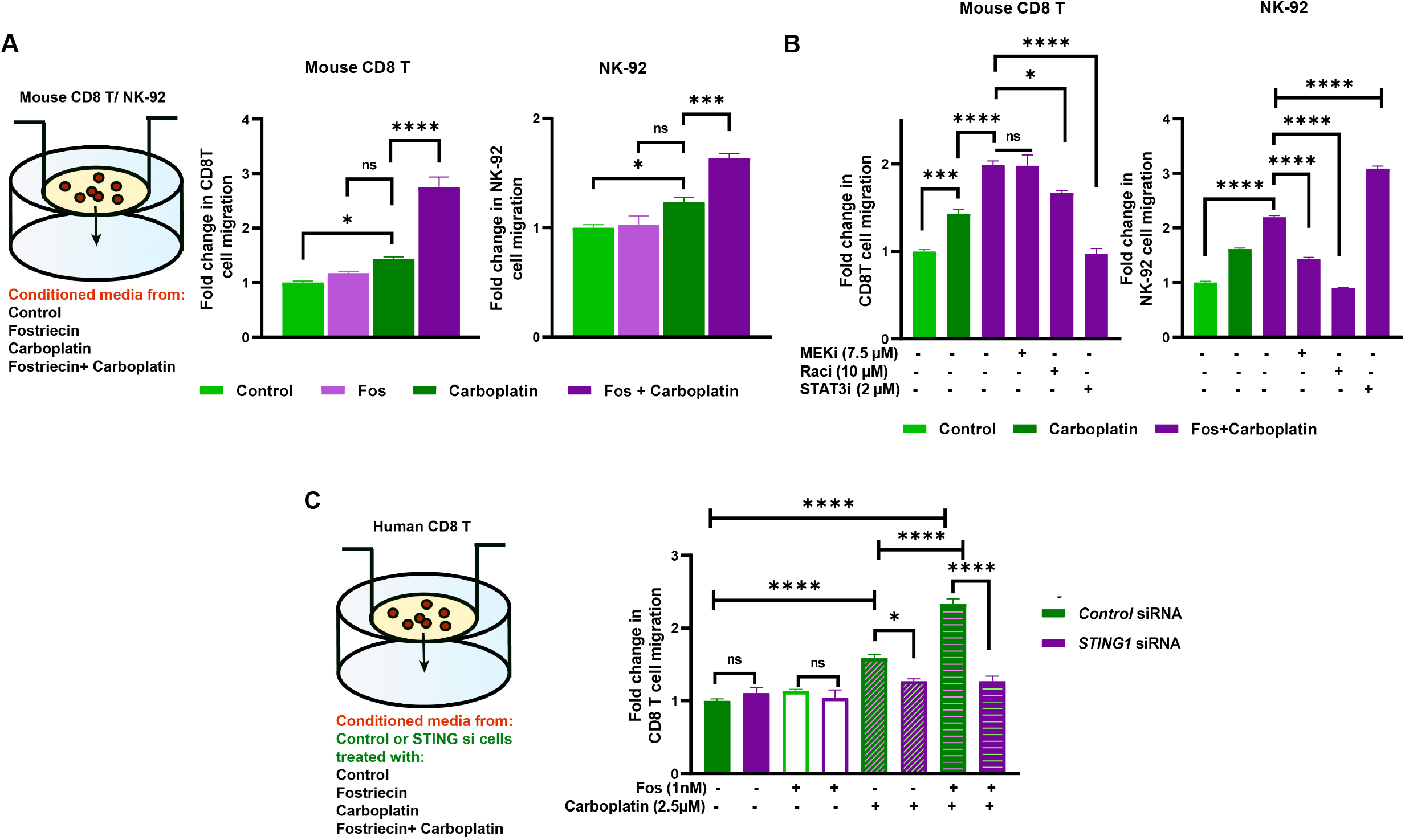
PP4 knockdown in OC cells enhances immune cell migration. (A) Schematic diagram of *in vitro* immune cell migration assay. Conditioned media was collected on day 5 from either ID8-p53KO (mouse) or OVCAR3 (human) cells ± Fos followed by carboplatin treatment at the indicated doses: 10 µM (ID8-p53KO) and 1 µM (OVCAR3). The conditioned media from each treatment group was loaded in the bottom chamber. Fold change in migration was calculated relative to control, mean ± SEM (n=3) is shown. *p<0.05, ***p<0.001 and ****p<0.0001. (B) Conditioned media was collected on day 5 from either ID8-p53KO or OVCAR3 cells were treated as described in previous experiment. In parallel, CD8 T or NK-92 cells were pre-treated with either MEKi (7.5 µM) or Raci (10 µM) or STAT3i (2 µM) for 15 min prior to adding in the transwell chamber. Conditioned media from each treatment group was loaded in the bottom chamber. Fold change in migration was calculated relative to control, mean ± SEM (n=3) is shown. *p<0.05, ***p<0.001 and ****p<0.0001. (C) Schematic diagram of *in vitro* CD8 T cell migration assay following *STING1* siRNA transfection in ovarian cancer cells. OVCAR8 cells were transfected with control or *STING1* siRNA ±Fos and carboplatin. On day 5, conditioned media was collected and used in the lower chamber as chemoattractant. Human CD8 T cells were isolated from the ascites of an ovarian cancer patient and used in the upper chamber. Fold change in migration (n=3) is shown, *p<0.05 and ****p<0.0001.

We next sought to determine whether blocking signaling downstream of chemokine and cytokine receptors could blunt the immune cell migration induced by the conditioned media from carboplatin and fostriecin treated OC cells. Consistent with the well-documented role of Rac signaling in chemokine-induced cell migration [34], we observed a significant reduction of both CD8 T and NK cell migration in the presence of a Rac inhibitor **(Figure 5B)**. However, we observed cell-type specific responses to MEK and STAT3 inhibitors. The STAT3 inhibitor, STATTIC, suppressed CD8 T cell migration but increased NK-92 cell migration **(Figure 5B)**. These results are consistent with previous reports as STAT3 signaling is known to be activated in migrating CD8 T cells in response to IL6 [35], whereas STAT3 is reported to have an opposing effect on NK cell migration [36]. Similar to STAT3 signaling, MEK inhibition also resulted in distinct cellular responses from CD8 T and NK cells. Interestingly, the MEK inhibitor suppressed NK cell migration, whereas CD8 T cell migration remained unaffected **(Figure 5B)**. To determine if cGAS-STING signaling played a role in the observed increase in T and NK cell migration, we knocked down *STING1* using siRNA in OVCAR8 cells **(Supplementary Figure 6B)**. Conditioned media from *STING1* knockdown OVCAR8 cells failed to stimulate T cell migration following PP4 inhibition **(Figure 5C)**. These data show that the pro-inflammatory signaling stimulated by PP4 inhibition is mediated by STING activation in OC cells.

### PP4C knockdown in tumor cells augmented NK cell activation and cytotoxicity against ovarian cancer

Since inhibition of PP4 activity led to increased inflammatory signaling and immune cell migration, next we examined whether PP4 subunit expression in OC correlates with immune cell infiltration using the Tumor Immune Estimation Resource (TIMER2.0). Gene expression deconvolution algorithms CIBERSORT (cell type identification by estimating the relative subset of known RNA transcripts) and xCELL were utilized to estimate the NK cell, CD8 T cell, and NKT cell subsets based on bulk RNAseq gene expression data from OC (TCGA). High transcript-level expression of *PPP4R3A* (SMEK1) and *PPP4R2* in OC correlated with a small but significant decrease in activated NK cell (CIBERSORT) and NKT (XCELL) infiltration. We observed a similar negative correlation in NK (p-value not significant) and NKT cell filtration (p-value significant) with high tumoral *PPP4R3B* expression. However, while we observed a negative trend with *PPP4C* expression, the adjusted p-value did not reach statistical significance using CIBERSORT **(Supplementary Figure 7)**. There was no significant correlation between expression of PP4 subunits and CD8 T cell infiltration (data not shown).

Studies have shown that OC is susceptible to NK cell-mediated killing and increased fractions of NK cells in ascites of OC patients correlate with improved outcomes [37, 38]. However, as we observed a significant negative correlation between RNA level expression of PP4 subunits and NK cell infiltration in OC, we next wanted to test our hypothesis that *PPP4C* or *PPP4R3B* knockdown can positively influence NK cell function against OC. We measured IFN-γ^+^ effector NK cells following co-culture with NK-92 cells and OVCAR8 transfected with either control, *PPP4C* or *PPP4R3B* siRNA. We saw a significant increase in IFN-γ^+^ NK cells in response to either *PPP4C* or *PPP4R3B* knockdown as compared to the control **(Figure 6A)**. Interestingly, while a further increase in the percentage of IFN-γ^+^ NK cell population was noted upon addition of carboplatin, it did not reach statistical significance over the siRNA alone group **(Figure 6A)**. To further determine if the increase in NK cell activation mediated by loss of PP4 contributed to an increase in NK cell-mediated OC cell killing, we co-cultured OC cell lines transfected with control, *PPP4C*, or *PPP4R3B* siRNA with NK-92 cells. Our results show that loss of *PPP4C* or *PPP4R3B* expression in OC enhances NK cell-directed OC cell killing **(Figure 6B)**. Carboplatin treatment induced a significant increase in killing with OVCAR8 cells, which was further increased in both *PPP4C* and *PPP4R3B* knockdown cells **(Figure 6B)**. In OVCAR3, even though there was increased killing in carboplatin treated cells over the siRNA alone group, it did not achieve statistical significance **(Supplementary Figure 8)**. We next assessed NK cell degranulation following a 3 h co-culture between NK-92 cells and OVCAR8 cells transfected with control, *PPP4C*, or *PPP4R3B* siRNA. We saw a significant increase in CD107a+ NK-92 cells in response to either *PPP4C* or *PPP4R3B* knockdown. We also observed a significant increase in CD107a+ NK-92 cell population upon co-culture with carboplatin treated cells (**Figure 6C**).

**Figure 6:**
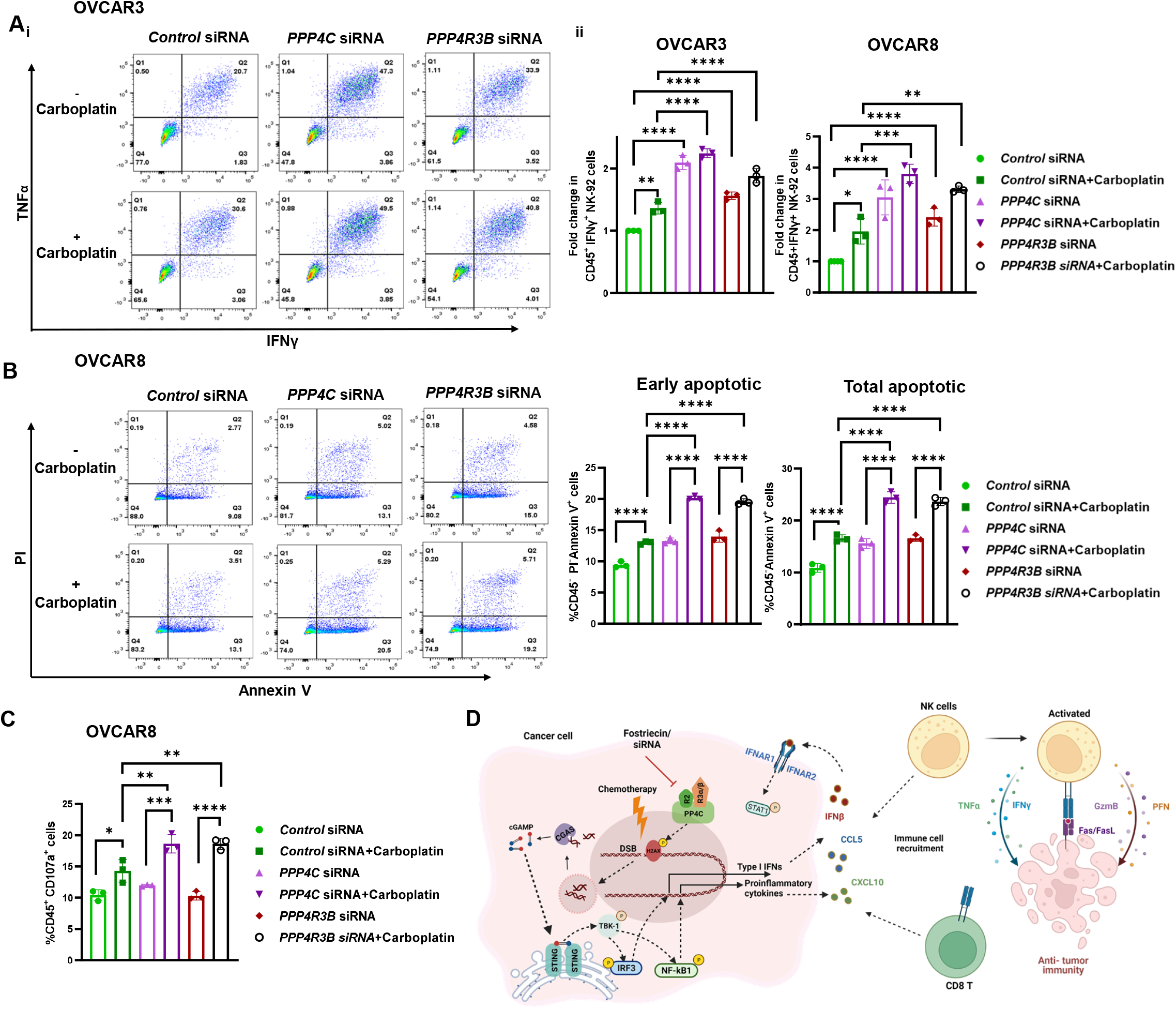
Loss of PP4 promotes NK cell activation and NK cell-directed OC killing. (A) OVCAR3 and OVCAR8 cells were transfected with either control or *PPP4C* or *PPP4R3B* siRNA, followed by carboplatin treatment at indicated doses: OVCAR3 (0.5 μM), OVCAR8 (2.5 μM). On day 5, cells were co-cultured with NK-92 (1:1) and immunostained for IFNγ and TNFα following PMA restimulation (i) Representative images of cytokine profile for OVCAR3 are shown. (ii) Relative fold changes in IFNγ+ cells were calculated for OVCAR3 (n=3) and OVCAR8 (n=3) and represented, mean± SD, *p<0.05, **p<0.01, ***p<0.001 and ****p<0.0001. (B) For NK-92 mediated cytotoxicity assays, OVCAR8 cells (n=3) transfected with control or *PPP4C* or *PPP4R3B* siRNA ± carboplatin treatment (2.5µM). NK-92 cells were added in 1:1 ratio. After 3 h, the cells were stained with Annexin V and PI and the CD45-negative cells were analyzed. Percentage of early and total apoptotic cells are shown for all treatment groups, mean± SD, ****p<0.0001. (C) Percentage of CD45^+^ CD107a^+^ cells following NK-92 and OVCAR8 co-culture is shown as bar graph, *p<0.05, **p<0.01, ***p<0.001 and ***p<0.0001.(D) Schematic diagram showing mechanism by which PP4C is involved in anti-tumor immunity. Loss of PP4, augments chemotherapy-induced DNA damage triggering a type I interferon response mediated by STING and STAT1 signaling. Increased chemokine production upon inhibition or knockdown of PP4C led to improved immune cell migration and NK cell-directed ovarian cancer cell death *in vitro*. Created with Biorender.com

## DISCUSSION

In this study, we provide direct evidence that loss of PP4 can augment inflammatory responses in OC (**Figure 6D**). Knockdown or pharmacological inhibition of PP4C activated type I interferon signaling leading to increased transcription of proinflammatory cytokines and STAT1 activation. Increased chemokines and cytokines from carboplatin treated, PP4-silenced OC cells enhanced immune cell migration. In addition, knockdown of either the catalytic subunit or regulatory subunit of PP4 in OC cells improved NK cell activation and NK cell-mediated OC killing.

PP4 plays a key role in DNA damage repair [10-13]. Dephosphorylation of DDR proteins by PP4 is critical for resolution of the DNA repair process [11]. PP4C and its regulatory subunits have previously been shown to be upregulated in human breast and lung tumors [39]. In this study, we found the PP4 subunits to be amplified and overexpressed in OC at the RNA level and were robustly expressed at the protein level in the majority of the OC cell lines tested. In a previous study, we had discovered that CT45 binds to and inhibits PP4, thereby enhancing carboplatin sensitivity [14]. We hypothesized that reducing PP4 activity via an inhibitor should generate a similar phenotype. Fostriecin, a commercially available phosphate ester produced by *Streptomyces pulveraceous*, was originally identified as an anti-tumor antibiotic that demonstrated anti-tumor function in xenograft mouse models [40]. Subsequently it was discovered that fostriecin was a potent, selective inhibitor of PP4 and PP2 phosphatases [41].

However, many of the effects of fostriecin, such as mitotic slippage, have more recently been attributed to its inhibition of PP4 rather than PP2A [42]. In addition, many OC tumors express high levels of the endogenous PP2A inhibitor, CIP2A [43], and therefore, PP2A activity is anticipated to be low in majority of OC tumors. As expected, fostriecin increased carboplatin chemosensitivity, an effect that was mimicked by the use of *PPP4C* siRNA.

Increased micronuclei formation as a result of DNA damage and chromosomal instability can lead to activation of the cGAS-STING pathway [44]. Cell-intrinsic cGAS/STING signaling leads to activation of type I inflammatory response and subsequent STAT1 activation [30]. We found that fostriecin combined with carboplatin led to increases in micronuclei formation, *IFNB1* transcript levels, and STAT1 activation in OC cells. The cGAS/STING pathway is known to activate downstream canonical NFκB [45] and we observed increased phosphorylation of p65 upon fostriecin treatment, as well as with PP4C knockdown in carboplatin treated OC cells. In addition, we found significant increases in the transcript levels of the pro-inflammatory cytokines and chemokines, *IL6, CCL5, CXCL10*, upon both PP4C knockdown and fostriecin treatment combined with carboplatin.

T helper 1 (T_H_1)-type chemokines, such as CXCL9 and CXCL10, have been shown to stimulate the recruitment of effector CD8 T cells and NK cells into the tumor, and these immune cells can modulate responses to ICB [33]. In OC, NK cells have been reported to co-infiltrate along with CD8 T cells and are strongly associated with patient survival [46]. Our results show that conditioned media collected from OC cells treated with fostriecin or *PPP4C* siRNA plus carboplatin, significantly increased both T and NK cell migration. Immune cell migration was found to be dependent on the STING pathway as STING knockdown in OC cells suppressed T cell migration in response to combination treatment with fostriecin and carboplatin. Additionally, we show that NK-92 cells co-cultured with PP4 knockdown OC cells had enhanced IFNγ levels, degranulation, and NK cell-mediated killing of OC cells. These data demonstrate a crucial role for tumorigenic PP4 in regulating NK cell function.

The introduction of immune checkpoint inhibitors into clinical care was a significant breakthrough for the field of cancer immunology. However, clinical trials have revealed that single agent ICB therapy is not effective in OC patients [47]. Recently, the combination of checkpoint blockade with PARP inhibitors has been shown to increase the objective response rate in OC [8]. Consequently, finding additional molecular targets involved in the DNA damage response that may influence ICB response is an area of immense translational interest. To our knowledge, this is the first report demonstrating that knockdown or pharmacologic inhibition of PP4 results in enhancement of immune cell effector function. Taken together, our results clearly identify PP4 as a potential target in the DDR pathway that could enhance the anti-tumor immune response in OC. In light of these findings, we believe PP4 to be an attractive therapeutic target in OC that warrants further investigation.

## Supporting information

Raja et al _Supplementary file_PP4

## Supplementary Figure Legend

**Supplementary Figure 1:** Genetic alterations of PP4 complex in ovarian cancer. (A) Oncoprint showing genetic and transcript level alterations of PP4 in ovarian cancer (n=202). The image was generated using cBioportal. (B) Comparison of RNA expression versus copy number for *PPP4R3A, PPP4R3B* and *PPP4R2* from pan-cancer TCGA data is plotted as bar graph, ****p<0.0001.(C) The transcript level expression of PPP4C along with proteomics score was obtained from Depmap portal and plotted as graph with RNA level expression on left -Y axis and proteomics score on right -Y axis. Proteomics score for PP4C was not available for OVCAR5 and 59M.

**Supplementary Figure 2:** Generation and p53 CRISPR knockout validation in ID8 mouse ovarian cancer cells. (A) Sequence of the target region of *TP53* gene from reference genome (black) non-targeting control, NTC (blue) and Trp53-/- clone E12e (green) are shown (top panel). gRNA sequences used to generate the clone are underlined. Sanger sequencing of the region around exon 5 of *TP53*-/- clone E12e and NTC are shown. The deleted region is highlighted in red and break points indicated in clone E12e (bottom panel). (B) Representative PCR of 538 base pair region spanning the targeted exon 5, showing deletion in ID8 E12e clone compared to NTC control. (C) Expression of p53 was determined in parental ID8 and three *TP53*-/- clones± carboplatin (100 µM) by immunoblot. Actin was used as loading control.

**Supplementary Figure 3:** Workflow for determining PP4 phosphatase activity is shown. Immunopurified PP4 complex in OVCAR3 and OVCAR4 cells were incubated with Fos (1nM) for 30 min. After substrate addition, relative fluorescence was measured and quantified. The results represented here are the average of 3 independent experiments ±SEM, *p < 0.05.

**Supplementary Figure 4:** Fostriecin augments carboplatin-induced DNA damage in OC cell lines. (A) Immunoblot analysis showing relative changes in γ-H2AX (S139) in OC cell lines treated ± Fos (1nM) and carboplatin treatment at 96 h. Fold changes in intensities are calculated over respective actin controls and indicated. The immunoblot densitometry was performed by ImageJ software (NIH), relative fold changes were calculated after normalization with corresponding β-actin bands. (B). Quantification of FANCD2 and γ-H2AX (S139) staining intensities (derived from the high throughput image analysis) were compared across different treatment groups. An average from 70-90 sites is represented as violin plots, p<0.0001.

**Supplementary Figure 5:** Loss of PP4 leads to increased inflammatory signaling and cytokine production in ID8-p53KO cells. (A) ID8-p53KO cells were treated ± Fos (1nM), followed by carboplatin (10µM). On day 5, cells were harvested and immunoblot analysis was performed to determine STAT1 phosphorylation levels across treatment groups. (B) ID8-p53KO cells were transfected with *PPP4C* siRNA followed by carboplatin (10µM). STAT1 phosphorylation is shown by western blot. (C) Transcript levels of *CCL5, CXCL10, IL6* and *IFNB1* following both Fos treatment and *PPP4C* siRNA transfection in ID8-p53KO were measured by quantitative PCR and relative fold change was calculated from 3 replicates is represented as bar graph. *p<0.05, * *p<0.01, ***p<0.001 and****p<0.0001.

**Supplementary Figure 6:** (A) Direct fostriecin treatment does not affect CD8 T cell activation. Splenocytes were harvested from OT I mice and stimulated with OVA peptide (SIINFEKL) in presence of varying doses of fostriecin. On Day 4, cells were restimulated with OVA in the presence of fostriecin and stained for intracellular TNFα and IFNγ. The percentage of cells expressing TNFα and IFNγ across treatment conditions is represented as bar graph (n=3). (B) Immunoblot showing STING1 protein levels in OVCAR8 cells transfected with either control or *STING1* siRNA and treated with Fos and carboplatin. Actin was used as loading control.

**Supplementary Figure 7**: PP4 subunit expression and immune cell infiltration. Correlation between the expression of PP4 subunits and infiltrating level of NK cells and NKT in ovarian cancer using CIBERSORT and xCELL on TIMER2.0 platform. Partial Rho and p values along with purity correction are shown.

**Supplementary Figure 8:** NK cell-mediated killing of HGSOC cells. OVCAR3 cells were transfected with control, *PPP4C* or *PPP4R3B* siRNA and treated ± carboplatin (2.5µM). Following co-culture, the cells were stained with Annexin V and PI and the CD45-negative cells were analyzed. Percentage of early and total apoptotic cells are shown for all treatment groups(n=4), ± SD, *p<0.05 and **p<0.01.

## Declarations

### Ethics approval and consent to participate

Ascites was collected from ovarian cancer patients undergoing primary debulking surgery by a gynecologic oncologist at the Mayo Clinic hospital, Arizona. Informed consent was obtained before surgery and the study was approved by the IRB of the Mayo Clinic. Collection of the OT-I T cells from mice was approved by the Institutional Animal Care and Use Committee of Mayo Clinic.

### Consent for publication

All authors have read the manuscript and have consented to publish.

### Availability of data and material

All data supporting the findings of this study are available within the article and its supplemental information files or obtained from the lead contact upon reasonable request.

## Author’s contributions

MC conceived and designed the study. RR contributed to the study design, performed experiments, and drafted the manuscript. RR, CW performed flow cytometry and CETSA. TR performed immunofluorescence staining and data analysis. EU contributed to manuscript writing. PM and KB provided resources. MC, PM, KB reviewed and edited the manuscript. MC supervised the work and is responsible for the overall content as guarantor.

## Acknowledgements

We are grateful to Scott Kaufmann for valuable advice and Larry Karnitz for providing the OVCAR8 DR-GFP cell line.

---

**Supplementary Table 1.**
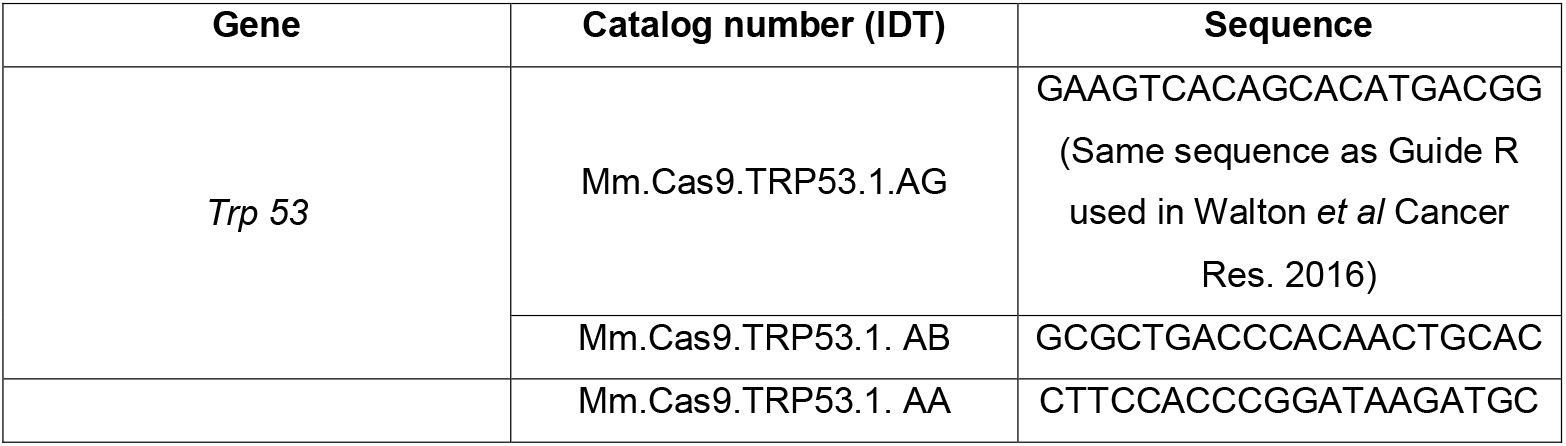
CRISPR crRNA sequences used in the study for mouse

**Supplementary Table 2.**
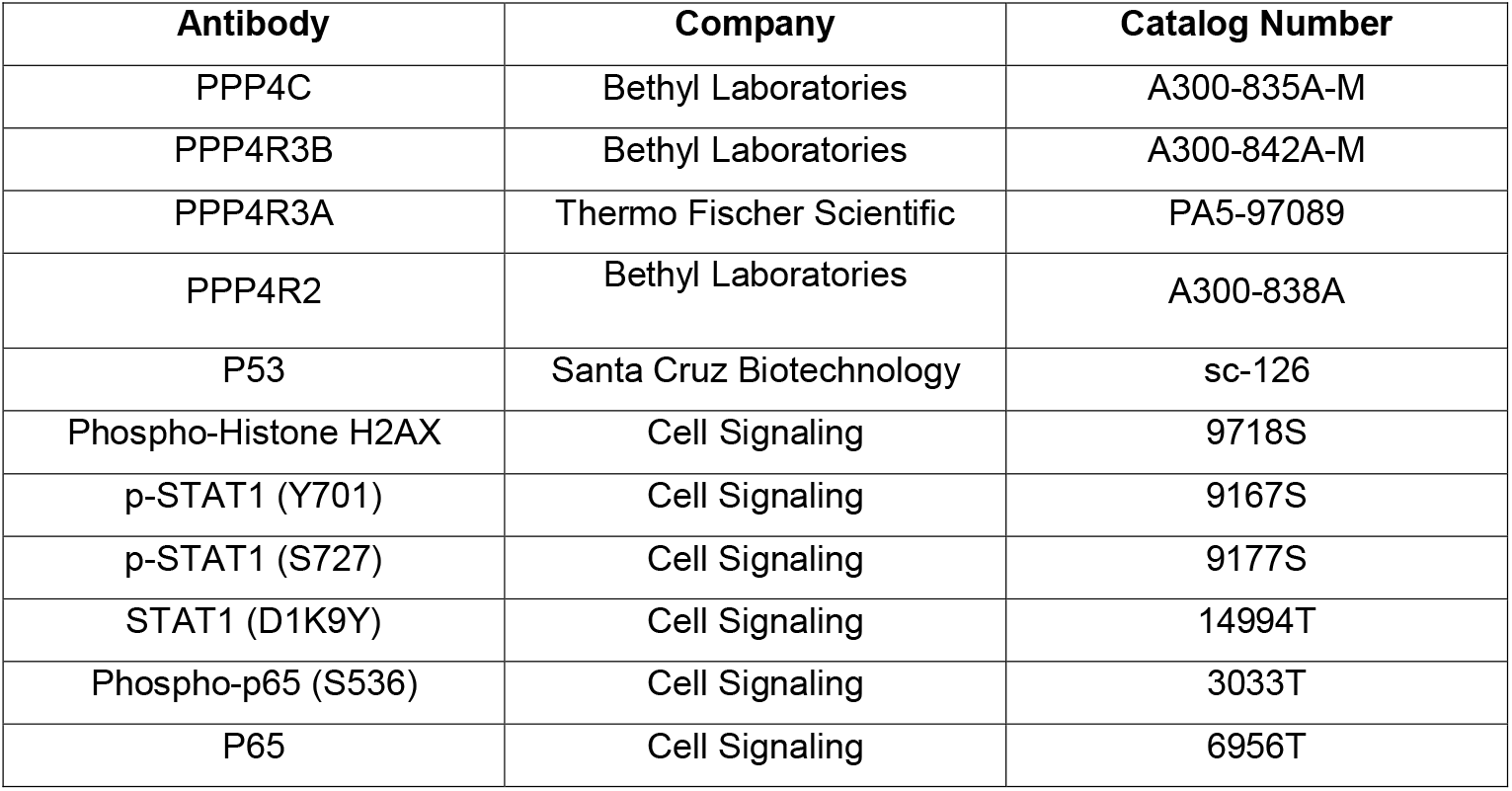
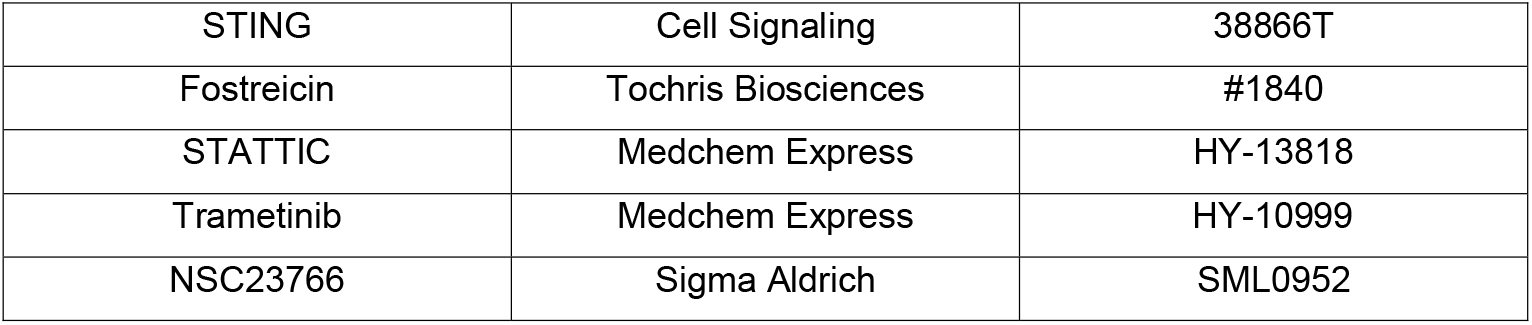
List of antibodies used for immunoblotting and inhibitors used in the study

**Supplementary Table 3.** List of Taqman probes used in the study

| Gene symbol | Company | Catalog Number: 4331182<br>Assay Number |
| --- | --- | --- |
| CXCL10 (human) | Thermo Fisher Scientific | Hs00171042_m1 |
| CXCL10 (mouse) | Thermo Fisher Scientific | Mm00445235_m1 |
| IL6 (human) | Thermo Fisher Scientific | Hs00174131_m1 |
| IL6 (mouse) | Thermo Fisher Scientific | Mm00446190_m1 |
| IFNB1 (human) | Thermo Fisher Scientific | Hs01077958_s1 |
| IFNB1 (mouse) | Thermo Fisher Scientific | Mm00439552_s1 |
| CCL5 (human) | Thermo Fisher Scientific | Hs00982282_m1 |
| CCL5 (mouse) | Thermo Fisher Scientific | Mm01302427_m1 |
| 18srRNA (human) | Thermo Fisher Scientific | Hs99999901_s1 |
| 18srRNA (mouse) | Thermo Fisher Scientific | Mm04277571_s1 |

**Supplementary Table 4.** List of antibodies used for flow cytometry in the study

| Antibody | Company | Catalog Number |
| --- | --- | --- |
| Ghost Dye UV 450 | Tonbo Biosciences | 13-0868-T500 |
| CD3 (anti-mouse) | Tonbo Biosciences | 70-0032-U500 |
| CD8a (anti-mouse) | Tonbo Biosciences | 75-0081-U100 |
| TNFA (anti-mouse) | BioLegend | 506308 |
| IFNG (anti-mouse) | Tonbo Biosciences | 50-7311-U100 |
| CD45 (ati-human) | Tonbo Biosciences | 25-0459-T100 |
| TNFA (anti-human) | BioLegend | 502932 |
| IFNG (anti-human) | BioLegend | 502509 |
| CD107a (anti-human) | BioLegend | 328640 |
| Annexin V | BD Biosciences | 556419 |

## REFERENCES

1. Siegel, R.L., et al., Cancer Statistics, 2021. CA Cancer J Clin, 2021. 71(1): p. 7–33.

2. Allemani, C., et al., Global surveillance of trends in cancer survival 2000-14 (CONCORD-3): analysis of individual records for 37 513 025 patients diagnosed with one of 18 cancers from 322 population-based registries in 71 countries. Lancet, 2018. 391(10125): p. 1023–1075.

3. Galluzzi, L., et al., The hallmarks of successful anticancer immunotherapy. Sci Transl Med, 2018. 10(459).

4. Zhang, L., et al., Intratumoral T cells, recurrence, and survival in epithelial ovarian cancer. New England Journal of Medicine, 2003. 348(3): p. 203–13.

5. Sato, E., et al., Intraepithelial CD8+ tumor-infiltrating lymphocytes and a high CD8+/regulatory T cell ratio are associated with favorable prognosis in ovarian cancer. Proc Natl Acad Sci U S A, 2005. 102(51): p. 18538–43.

6. Kandalaft, L.E., K. Odunsi, and G. Coukos, Immunotherapy in Ovarian Cancer: Are We There Yet? J Clin Oncol, 2019. 37(27): p. 2460–2471.

7. Ding, L., et al., PARP Inhibition Elicits STING-Dependent Antitumor Immunity in Brca1-Deficient Ovarian Cancer. Cell Reports, 2018. 25(11): p. 2972–2980.e5.

8. Konstantinopoulos, P.A., et al., Single-Arm Phases 1 and 2 Trial of Niraparib in Combination With Pembrolizumab in Patients With Recurrent Platinum-Resistant Ovarian Carcinoma. JAMA Oncology, 2019. 5(8): p. 1141–1149.

9. Park, J. and D.H. Lee, Functional roles of protein phosphatase 4 in multiple aspects of cellular physiology: a friend and a foe. BMB Rep, 2020. 53(4): p. 181–190.

10. Lee, D.H., et al., A PP4 phosphatase complex dephosphorylates RPA2 to facilitate DNA repair via homologous recombination. Nat Struct Mol Biol, 2010. 17(3): p. 365–72.

11. Chowdhury, D., et al., A PP4-phosphatase complex dephosphorylates gamma-H2AX generated during DNA replication. Mol Cell, 2008. 31(1): p. 33–46.

12. Lee, D.H., et al., Dephosphorylation enables the recruitment of 53BP1 to double-strand DNA breaks. Mol Cell, 2014. 54(3): p. 512–25.

13. Lee, D.H., et al., Phosphoproteomic analysis reveals that PP4 dephosphorylates KAP-1 impacting the DNA damage response. The EMBO Journal, 2012. 31(10): p. 2403–2415.

14. Coscia, F., et al., Multi-level Proteomics Identifies CT45 as a Chemosensitivity Mediator and Immunotherapy Target in Ovarian Cancer. Cell, 2018. 175(1): p. 159–170.e16.

15. Walton, J., et al., CRISPR/Cas9-Mediated Trp53 and Brca2 Knockout to Generate Improved Murine Models of Ovarian High-Grade Serous Carcinoma. Cancer Res, 2016. 76(20): p. 6118–6129.

16. Cerami, E., et al., The cBio cancer genomics portal: an open platform for exploring multidimensional cancer genomics data. Cancer Discov, 2012. 2(5): p. 401–4.

17. Cline, M.S., et al., Exploring TCGA Pan-Cancer data at the UCSC Cancer Genomics Browser. Sci Rep, 2013. 3: p. 2652.

18. Li, T., et al., TIMER2.0 for analysis of tumor-infiltrating immune cells. Nucleic Acids Res, 2020. 48(W1): p. W509–W514.

19. Newman, A.M., et al., Robust enumeration of cell subsets from tissue expression profiles. Nat Methods, 2015. 12(5): p. 453–7.

20. Sturm, G., et al., Comprehensive evaluation of transcriptome-based cell-type quantification methods for immuno-oncology. Bioinformatics, 2019. 35(14): p. i436–i445.

21. Guzman, C., et al., ColonyArea: an ImageJ plugin to automatically quantify colony formation in clonogenic assays. PLoS One, 2014. 9(3): p. e92444.

22. Jafari, R., et al., The cellular thermal shift assay for evaluating drug target interactions in cells. Nature Protocols, 2014. 9(9): p. 2100–2122.

23. Curtis, M., et al., Fibroblasts Mobilize Tumor Cell Glycogen to Promote Proliferation and Metastasis. Cell Metabolism, 2018. 29(1): p. 141–155.e9.

24. Kanakkanthara, A., et al., ZC3H18 specifically binds and activates the BRCA1 promoter to facilitate homologous recombination in ovarian cancer. Nat Commun, 2019. 10(1): p. 4632.

25. Galeano Nino, J.L., et al., Cytotoxic T cells swarm by homotypic chemokine signalling. Elife, 2020. 9.

26. Chava, S., et al., Measurement of Natural Killer Cell-Mediated Cytotoxicity and Migration in the Context of Hepatic Tumor Cells. J Vis Exp, 2020(156).

27. Lopez-Martinez, D., C.C. Liang, and M.A. Cohn, Cellular response to DNA interstrand crosslinks: the Fanconi anemia pathway. Cell Mol Life Sci, 2016. 73(16): p. 3097–114.

28. Motwani, M., S. Pesiridis, and K.A. Fitzgerald, DNA sensing by the cGAS–STING pathway in health and disease. Nature Reviews Genetics, 2019. 20(11): p. 657–674.

29. Meissl, K., et al., The good and the bad faces of STAT1 in solid tumours. Cytokine, 2017. 89: p. 12–20.

30. Harding, S.M., et al., Mitotic progression following DNA damage enables pattern recognition within micronuclei. Nature, 2017. 548(7668): p. 466–470.

31. Abe, T. and G.N. Barber, Cytosolic-DNA-mediated, STING-dependent proinflammatory gene induction necessitates canonical NF-kappaB activation through TBK1. J Virol, 2014. 88(10): p. 5328–41.

32. Wennerberg, E., et al., CXCL10-induced migration of adoptively transferred human natural killer cells toward solid tumors causes regression of tumor growth in vivo. Cancer Immunol Immunother, 2015. 64(2): p. 225–35.

33. Nagarsheth, N., M.S. Wicha, and W. Zou, Chemokines in the cancer microenvironment and their relevance in cancer immunotherapy. Nature Reviews Immunology, 2017. 17(9): p. 559–572.

34. Steffen, A., et al., Rac function is crucial for cell migration but is not required for spreading and focal adhesion formation. J Cell Sci, 2013. 126(Pt 20): p. 4572–88.

35. McLoughlin, R.M., et al., IL-6 trans-signaling via STAT3 directs T cell infiltration in acute inflammation. Proc Natl Acad Sci U S A, 2005. 102(27): p. 9589–94.

36. Cacalano, N.A., Regulation of Natural Killer Cell Function by STAT3. Front Immunol, 2016. 7: p. 128.

37. Hoogstad-van Evert, J.S., et al., Peritoneal NK cells are responsive to IL-15 and percentages are correlated with outcome in advanced ovarian cancer patients. Oncotarget, 2018. 9(78): p. 34810–34820.

38. Hoogstad-van Evert, J.S., et al., Harnessing natural killer cells for the treatment of ovarian cancer. Gynecol Oncol, 2020. 157(3): p. 810–816.

39. Wang, B., et al., Protein phosphatase PP4 is overexpressed in human breast and lung tumors. Cell Res, 2008. 18(9): p. 974–7.

40. Wilbur R. Leopold, J.L.S., Amie E. Mertus, James M. Nelson, Billy J. Roberts, and Robert C. Jackson, Anticancer Activity of the Structurally Novel Antibiotic CI-920 and Its Analogues1. Cancer Research, 1984. 44: p. 1928–1934.

41. Lewy, D.S., et al., Fostriecin: Chemistry and Biology. Current Medicinal Chemistry, 2002. 9(22): p. 2005–2032.

42. Theobald, B., et al., Suppression of Ser/Thr Phosphatase 4 (PP4C/PPP4C) Mimics a Novel Post-Mitotic Action of Fostriecin, Producing Mitotic Slippage Followed by Tetraploid Cell Death. Molecular Cancer Research, 2013. 11(8): p. 845–855.

43. Böckelman, C., et al., Prognostic role of CIP2A expression in serous ovarian cancer. British Journal of Cancer, 2011. 105(7): p. 989–995.

44. Sen, T., et al., Targeting DNA Damage Response Promotes Antitumor Immunity through STING-Mediated T-cell Activation in Small Cell Lung Cancer. Cancer Discovery, 2019. 9(5): p. 646–661.

45. Decout, A., et al., The cGAS–STING pathway as a therapeutic target in inflammatory diseases. Nature Reviews Immunology, 2021. 21(9): p. 548–569.

46. Webb, J.R., et al., Tumor-infiltrating lymphocytes expressing the tissue resident memory marker CD103 are associated with increased survival in high-grade serous ovarian cancer. Clin Cancer Res, 2014. 20(2): p. 434–44.

47. Varga, A., et al., Pembrolizumab in patients with programmed death ligand 1-positive advanced ovarian cancer: Analysis of KEYNOTE-028. Gynecol Oncol, 2019. 152(2): p. 243–250.

