## Supplementary material for "PP4 inhibition sensitizes ovarian cancer to NK cell-mediated cytotoxicity via STAT1 activation and inflammatory signaling": Raja et al _Supplementary file_PP4

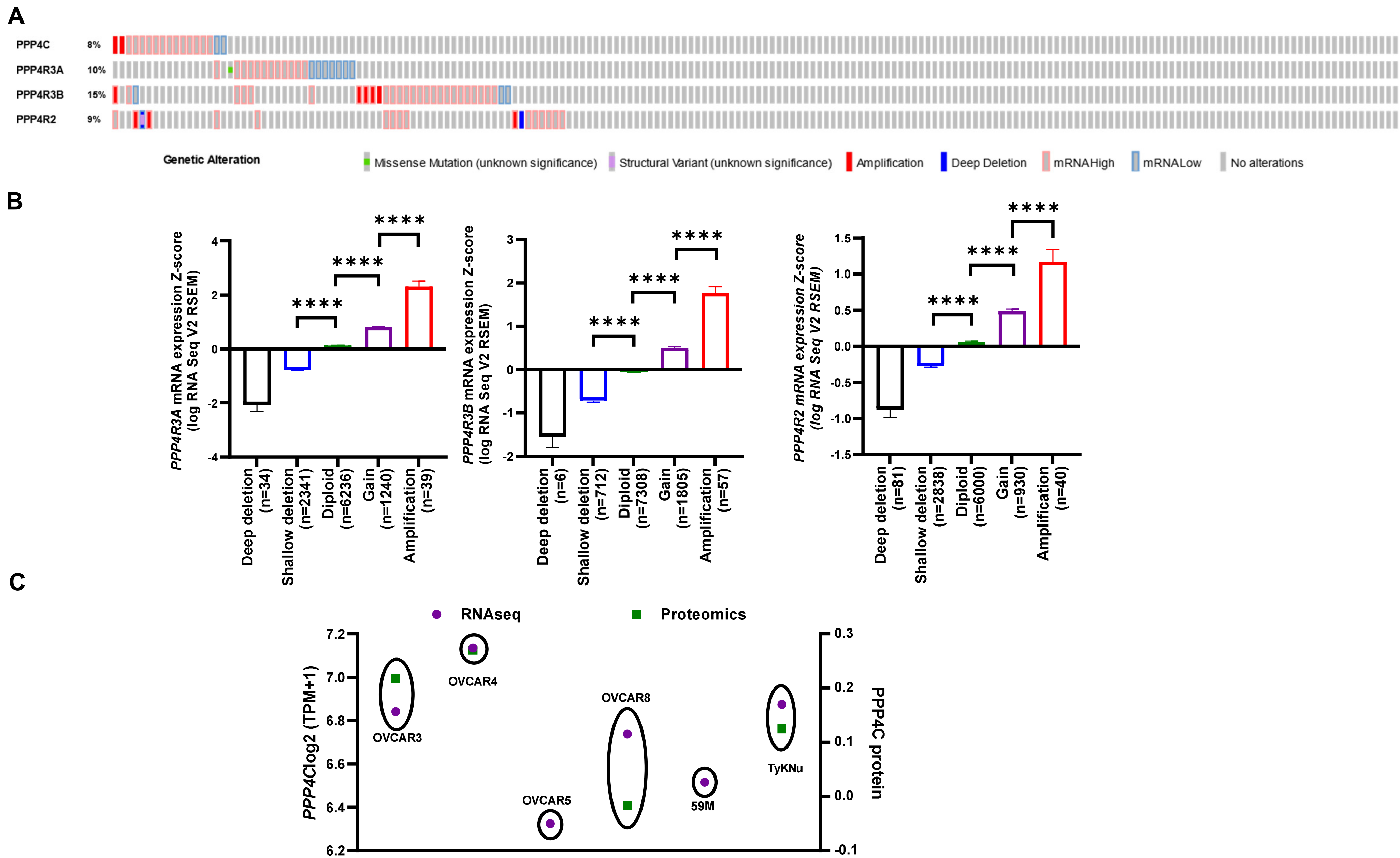


**Supplementary Fig. 1**

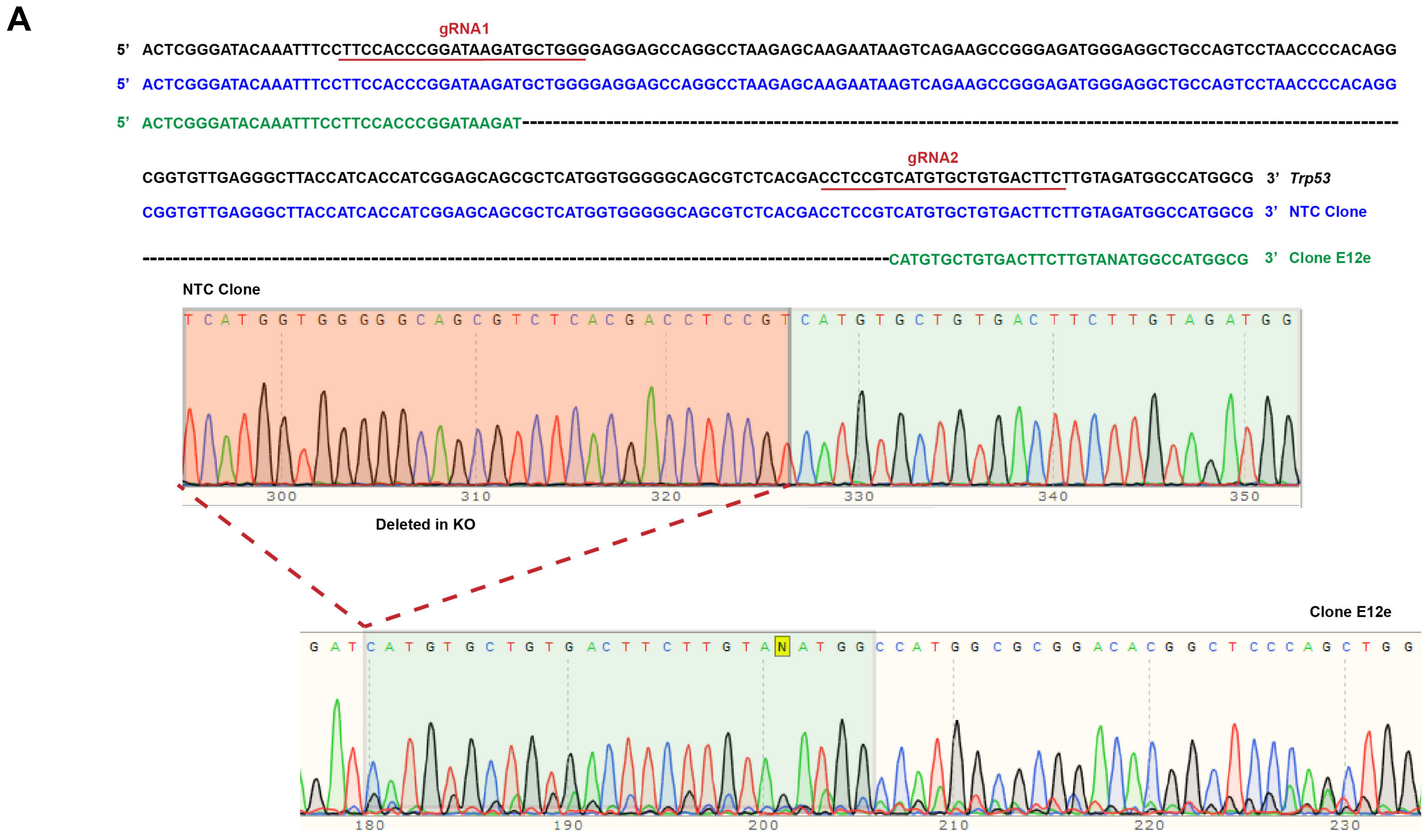

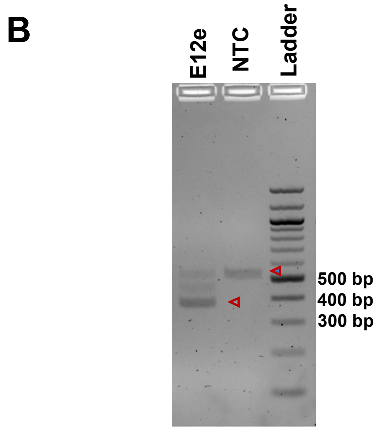

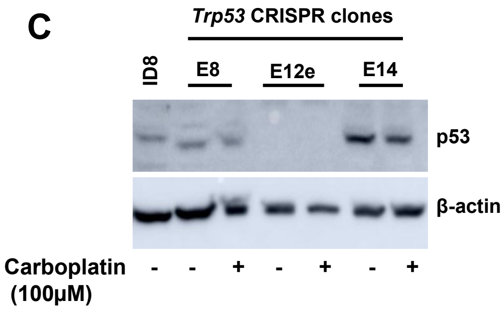


**Supplementary Fig. 2**

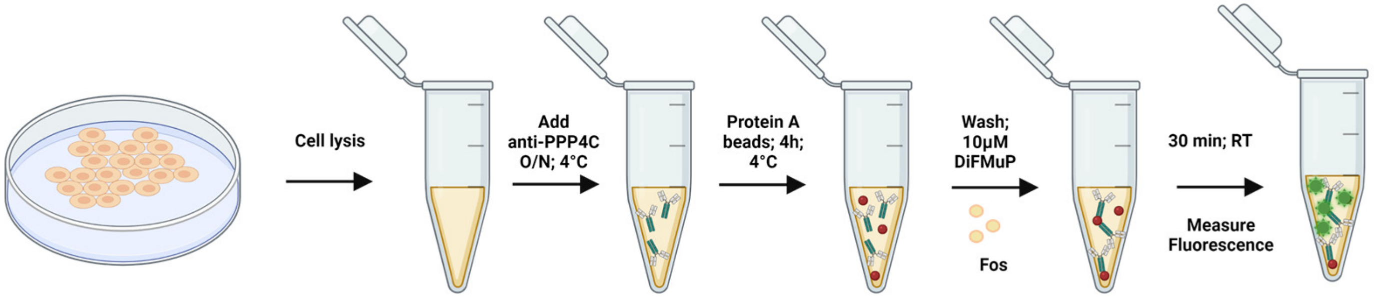


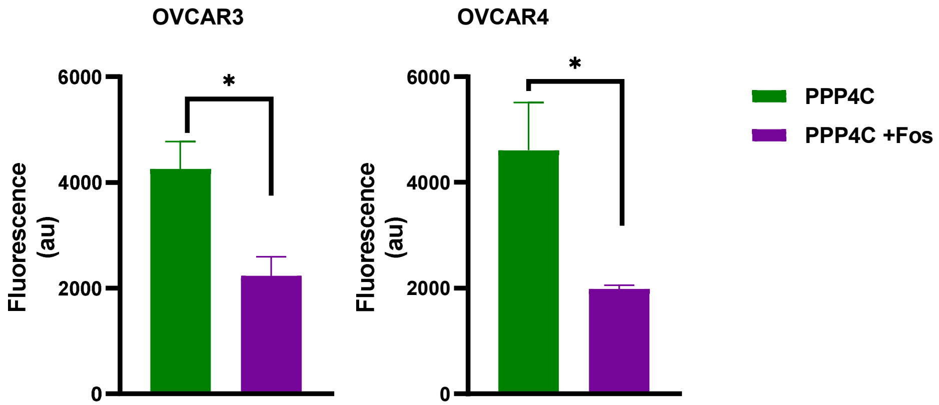


**Supplementary Fig. 3**

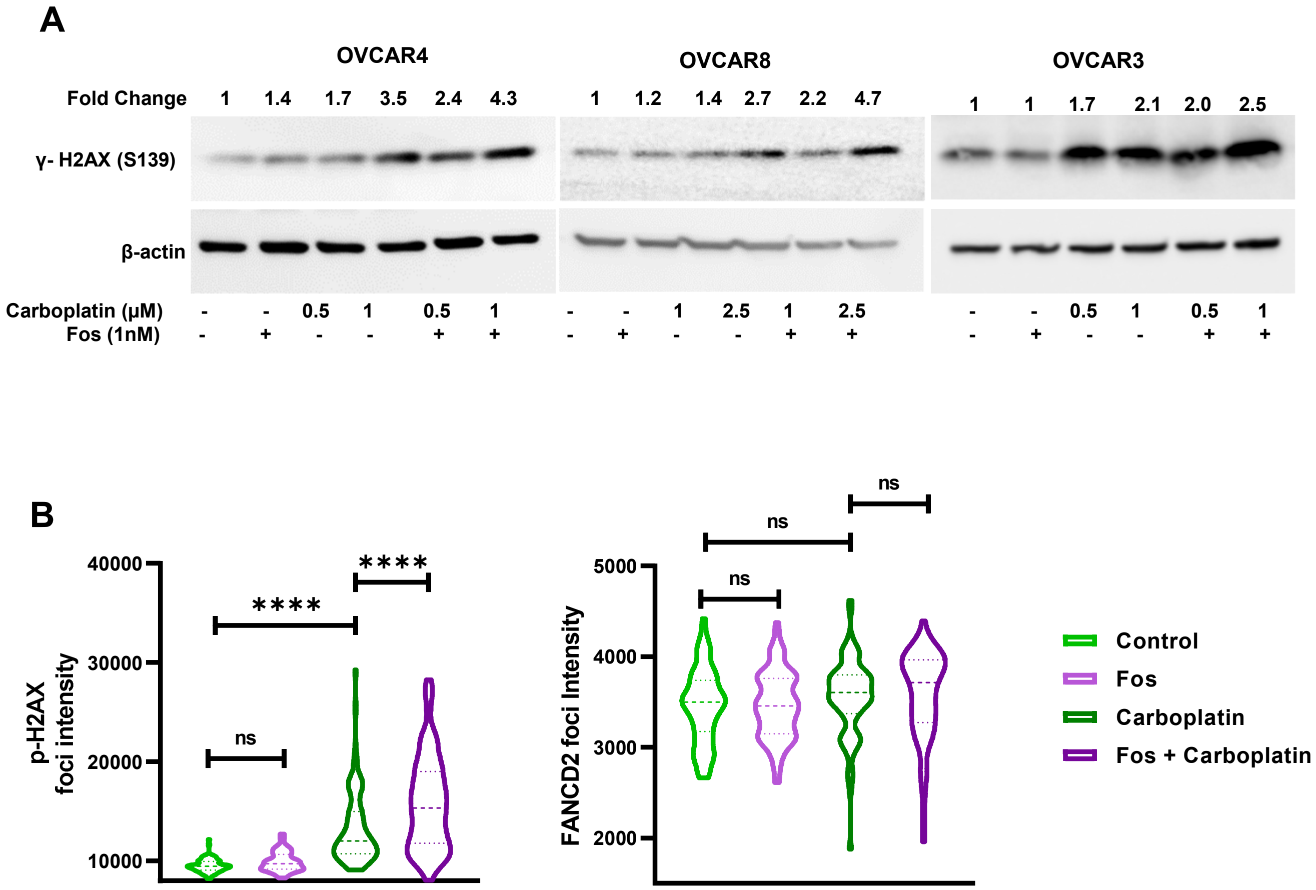


**Supplementary Fig. 4**

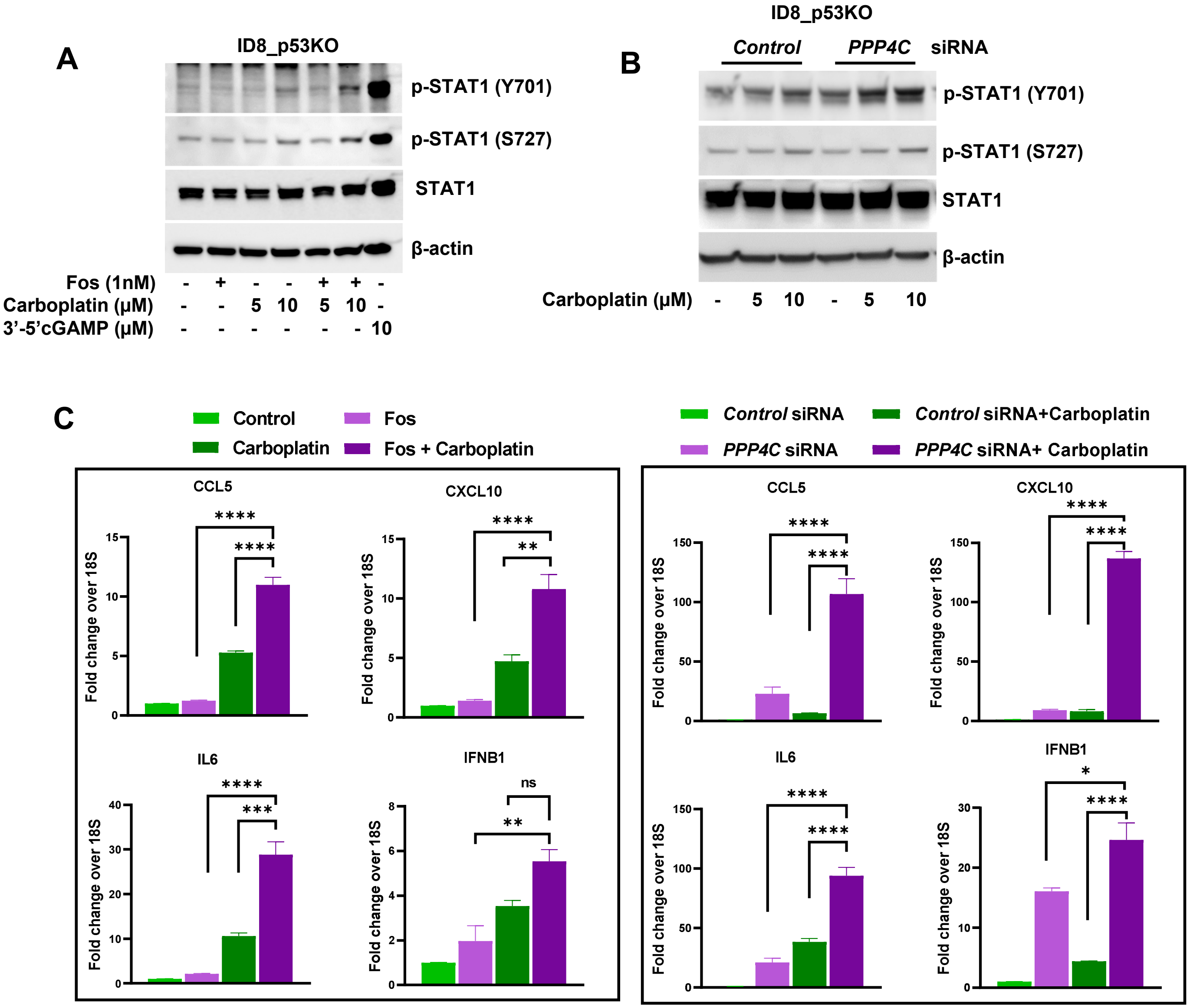


Supplementary Fig. 5

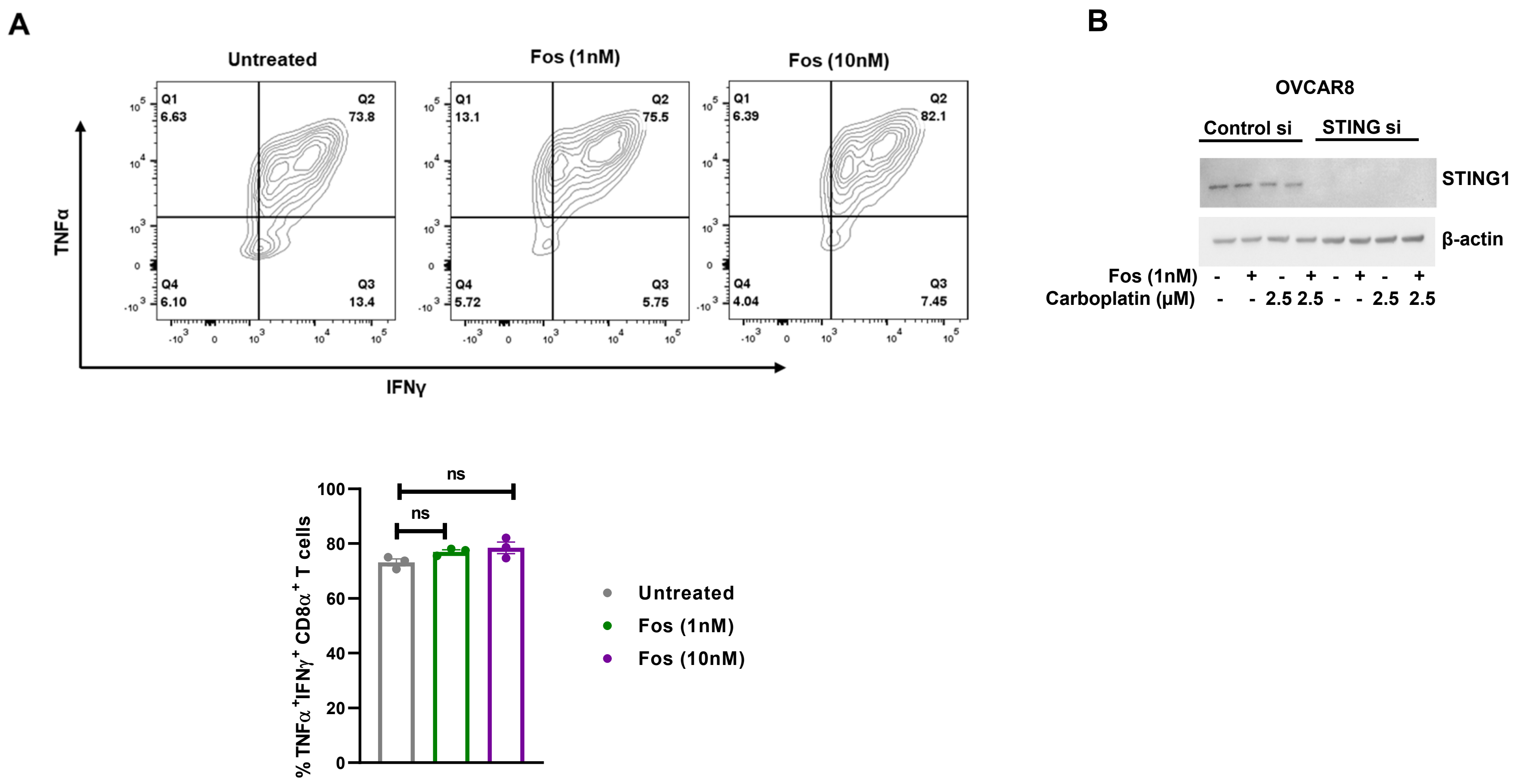


**Supplementary Fig. 6**

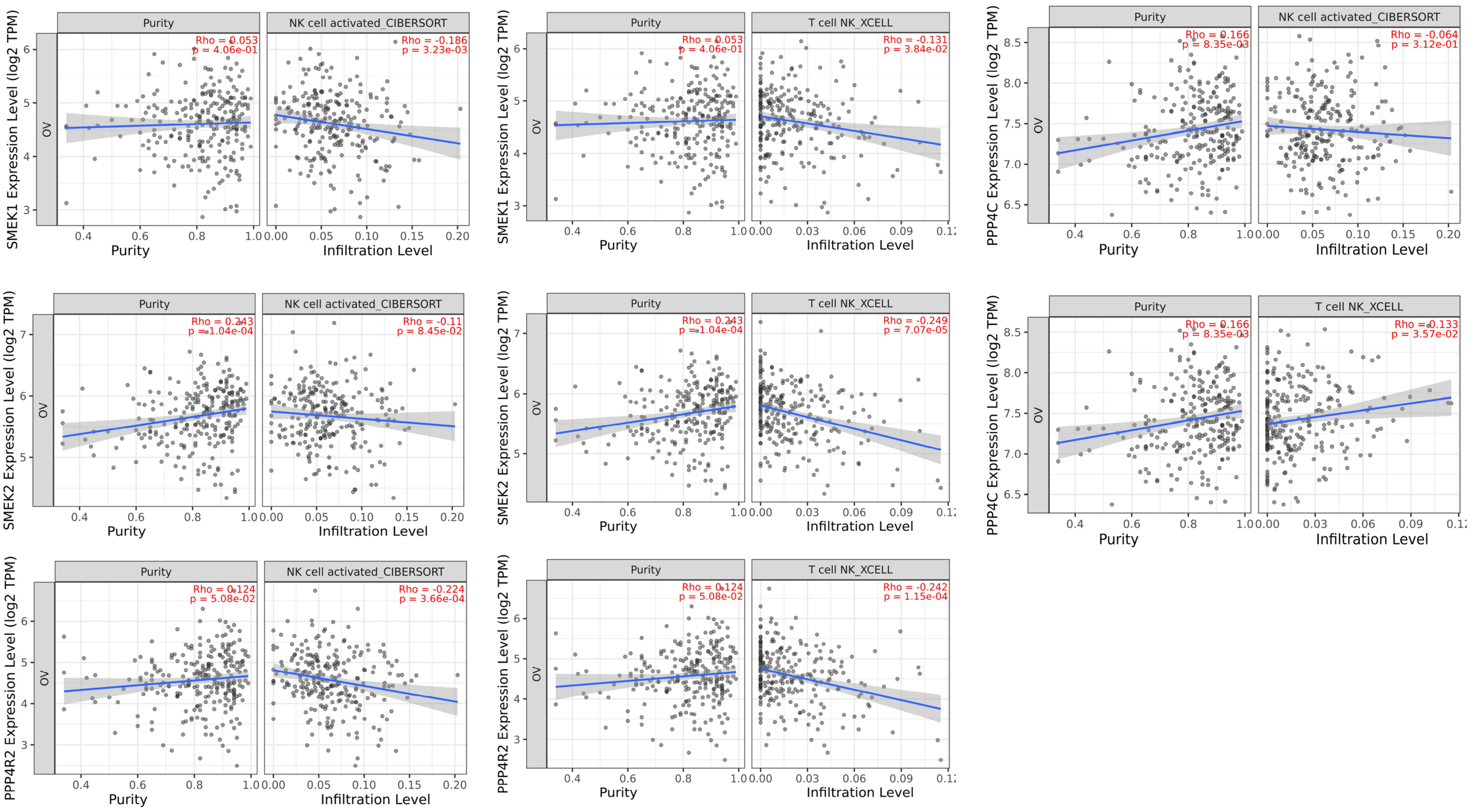


**Supplementary Fig. 7**

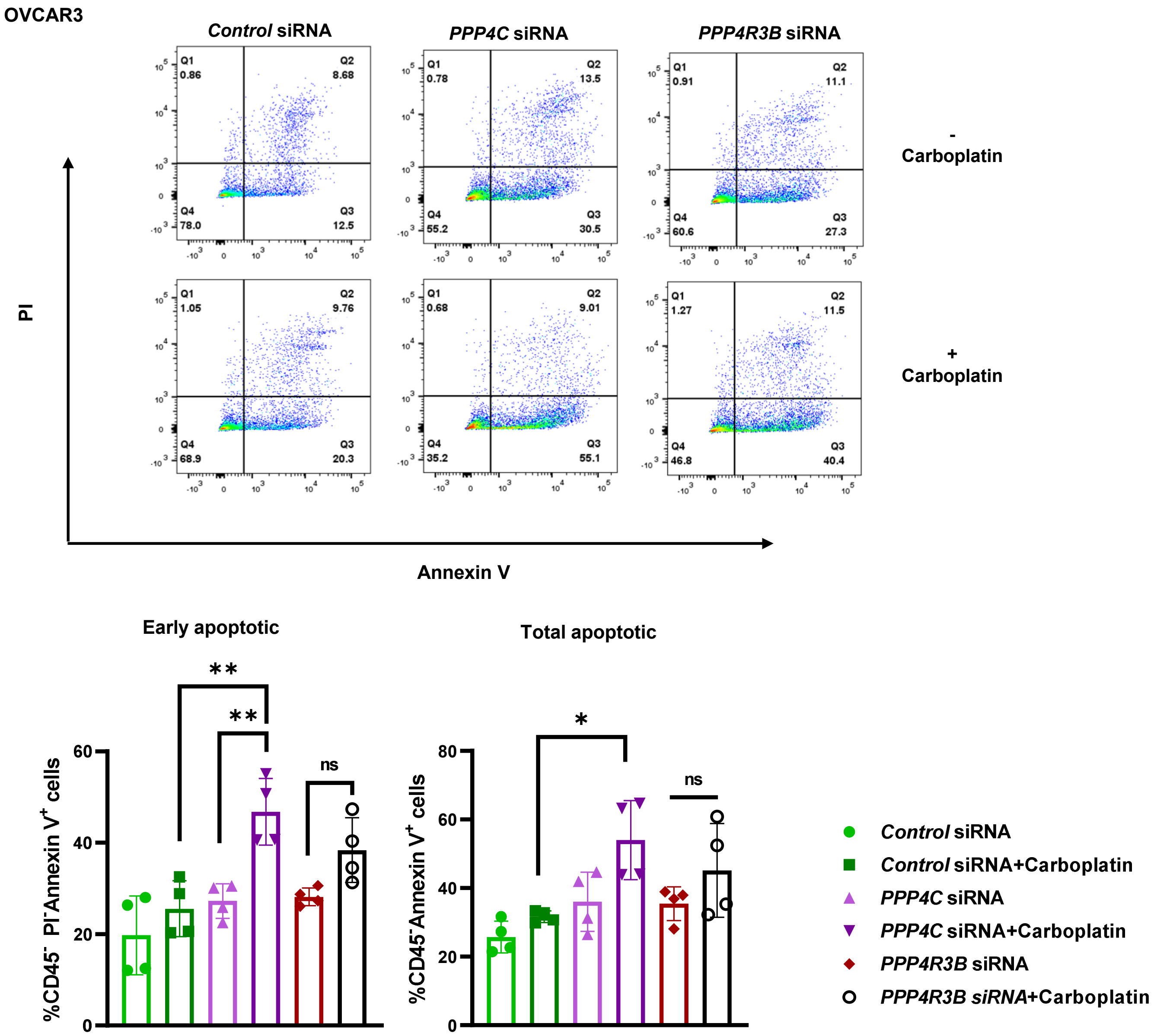


**Supplementary Fig. 8**
